# Spatiotemporal modeling reveals geometric dependence of AMPAR dynamics on dendritic spine morphology

**DOI:** 10.1101/2022.05.31.494202

**Authors:** M. K. Bell, C. T. Lee, P. Rangamani

**Author notes:** Corresponding Author | |.

## Abstract

The modification of neural circuits depends on the strengthening and weakening of synaptic connections. Synaptic strength is often correlated to the density of the ionotropic, glutamateric receptors, AMPAR, (*α*-amino-3-hydroxy-5-methyl-4-isoxazolepropionic acid receptor) at the postsynaptic density (PSD). While AMPAR density is known to change based on complex biological signaling cascades, the effect of geometric factors such as dendritic spine shape, size, and curvature remain poorly understood. In this work, we developed a deterministic, spatiotemporal model to study the dynamics of AMPAR during long term potentiation (LTP). This model includes a minimal set of biochemical events that represent the upstream signaling events, trafficking of AMPAR to and from the PSD, lateral diffusion in the plane of the spine membrane, and the presence of an extrasynaptic AMPAR pool. Using idealized and realistic spine geometries, we show that the dynamics and increase of bound AMPAR at the PSD depends on a combination of endo- and exocytosis, membrane diffusion, availability of free AMPAR, and intracellular signaling interactions. We also found non-monotonic relationships between spine volume and change in AMPAR at the PSD, suggesting that spines restrict changes in AMPAR to optimize resources and prevent runaway potentiation.

**Significance Statement:** Synaptic plasticity involves dynamic biochemical and physical remodeling of small protrusions called dendritic spines along the dendrites of neurons. Proper synaptic functionality within these spines requires changes in receptor number at the synapse, which has implications for down-stream neural functions, such as learning and memory formation. In addition to being signaling subcompartments, spines also have unique morphological features that can play a role in regulating receptor dynamics on the synaptic surface. We have developed a spatiotemporal model that couples biochemical signaling and receptor trafficking modalities in idealized and realistic spine geometries to investigate the role of biochemical and biophysical factors in synaptic plasticity. Using this model, we highlight the importance of spine size and shape in regulating bound AMPAR dynamics that govern synaptic plasticity, and predict how spine shape might act to reset synaptic plasticity as a built-in resource optimization and regulation tool.

## Introduction

Dendritic spines are small protrusions along dendrites that serve as signaling subcompartments and have characteristic shapes associated with their development state, learning, memory formation, and disease states [1, 2]. Most excitatory synapses are housed in dendritic spines and regulate the strength of their connections through biochemical and structural modifications in a process knkown as synaptic plasticity. In particular, the number of AMPARs, *α*-amino-3-hydroxy-5-methyl-4-isoxazolepropionic acid receptors, at the postsynaptic density (PSD) modulates the sensitivity of the synapse to presynaptic glutamate release, effectively tuning the synaptic connection strength [3, 4]. Since AMPAR is an important indicator of synaptic plasticity, specifically long-term potentiation (LTP) and long-term depression (LTD), it is vital to understand the role played by different factors to regulate AMPAR number density [5, 6].

The factors that influence AMPAR can be classified into biophysical and biochemical features. Key among these factors are the (a) upstream signaling network [7], (b) trafficking mechanisms and their location specificity [8–10], (c) lateral diffusion in the plane of the spine membrane [11, 12], and (d) size and shape of the spine [13]. We briefly summarize the key experimental observations and open questions in the literature with respect to AMPAR dynamics at the PSD with a specific focus on LTP.

- **AMPAR density increases during LTP** AMPAR interacts with a variety of membrane bound and cytosolic proteins [14]. In particular, it binds to PSD-95 (SAP90) at the PSD where it colocalizes into clusters [15–18]. AMPAR density at the PSD is used as a readout for synaptic plasticity, with long term potentiation (LTP) associated with an increase in AMPAR bound to PSD95 at the synapse [19]. During LTP induction, AMPAR is expected to increase up to 200% at the PSD within a few minutes [1, 9, 20, 21].
- **Role of CaMKII in AMPAR dynamics in LTP** CaMKII is an abundant, vital protein in spines. It interacts with AMPAR, along with a plethora of other species in dendritic spines. CaMKII is implicated in controlling the level of AMPAR on spine membranes through exocytosis rates [22], various binding/signaling interactions [23], and by triggering AMPAR influx into the spine [24, 25]. While CaMKII is known to autophosphorylate, its exact dynamics after phosphorylation have been debated as either a bistable switch with sustained elevated activity (bistable model) [26] or exponentially decaying over time (monostable model) [27, 28]. Since CaMKII is a key signaling species underlying synaptic plasticity, its spatiotemporal dynamics are of particular interest as they can significantly influence downstream AMPAR dynamics [29–33].
- **Role of spine size and shape** Dendritic spines have characteristic shapes and sizes related to their developmental stage, activation history, and function [34, 35]. Spines are typically categorized into four subtypes - filopodial, stubby, thin, and mushroom. Thin and mushroom spines tend to be more prevalent in the adult brain, where mushroom spines tend to be larger and more stable compared to the smaller and more adaptive thin spines [36, 37]. It is experimentally observed that PSD size correlations to spine volume [38], and AMPAR number at the PSD is proportional to synaptic area and spine head volume [39–41]. However, more recent analysis have brought these exact relationships under question suggesting a more complex regulation of AMPAR at the PSD [42]. Furthermore, the shape of dendritic spines is known to influence signaling [43–47], but it is unknown how spine geometry couples with trafficking mechanisms and signaling to regulate AMPAR dynamics at the PSD.
- **Role of trafficking** AMPARs are packaged in endosomes inside the cytoplasm which can exchange with the membrane via endocytosis and exocytosis [9, 10]. Kinase and phosphatase activity appear to influence these rates, with CaMKII and PP1 dynamics leading to exocytosis and endocytosis, respectively [14, 48, 49]. The exact location of endo/exocytosis remains unclear, although there is evidence indicating that these processes occur near the PSD, on the dendrite, or at various sites regulated by synaptic activity [50, 51].
- **Role of lateral diffusion and extrasynaptic pool** AMPAR undergoes lateral membrane diffusion at different rates in various regions of the spine membrane, depending on its biochemical interactions (estimates of 0.04-0.059 µm^2^ s^−1^ when bound to stargazin outside of the PSD; 0.007 µm^2^ s^−1^ when bound to PSD95 at the PSD) [11, 12, 52, 53]. Extrasynaptic pools of AMPAR on the membrane have been observed on dendritic spines and along the dendrite [54]. It has been proposed that CaMKII signaling can influence the trafficking of AMPAR from these pools into activated dendritic spines [22–24].

While these different aspects have been studied experimentally [15, 50, 54–57] and computationally [6, 7, 29, 48, 58, 59], a comprehensive spatiotemporal model that accounts for these different factors is missing in the literature. The roles of spatial localization and membrane-cytosol crosstalk of these biochemical interactions and the role of dendritic spine morphology in these processes are not well-known. The bulk-surface coupling between the cytosol and the synaptic membrane is likely a crucial mediator of AMPAR density [45, 60–62]. Additionally, surface diffusion is also an important parameter for governing the receptor dynamics [63–66]. Here, we developed a spatiotemporal model to investigate how spine morphology, trafficking, and signaling interact to regulate AMPAR density at the PSD. We sought to answer the following specific questions: Does spine morphology influence different underlying biochemical signaling pathways that determine AMPAR dynamics? How do geometric features affect trafficking modalities and therefore AMPAR dynamics? And finally, how do signaling dynamics, spine morphology, and trafficking modalities couple to regulate AMPAR density at the PSD in realistic spine geometries?

To answer these questions, we leverage a set of compartmental models of AMPAR dynamics and convert into a spatial, multi-compartment reaction-diffusion model. In a companion paper, we developed a compartmental model of AMPAR dynamics where we explored two different CaMKII dynamics that arise from multistability [29]. Expanding on that study, in this work, we found that i) both CaMKII models produce sustained but different elevated steady state bound AMPAR levels, ii) bound AMPAR temporal dynamics show three key timescales associated with the different trafficking mechanisms, and iii) both idealized and real spine geometries show a positive correlation between total receptor number at the PSD and spine volume to surface area ratio. They also exhibit a size-dependent resetting in the percent change of AMPAR at the PSD. This could act to optimize resources and prevent excessive synaptic plasticity in spines of certain sizes.

### Model Development

To investigate AMPAR spatiotemporal dynamics, we construct a simplified biochemical signaling network describing AMPAR dynamics, on the timescale of minutes, in hippocampal pyramidal CA1 neurons using deterministic reaction-diffusion equations, see Fig. 1a and [29]. Here we list the model assumptions, describe the key steps in the biochemical signaling cascade, provide the governing equations, and describe how the idealized and realistic spine morphologies were used for simulation purposes.

**Figure 1:**
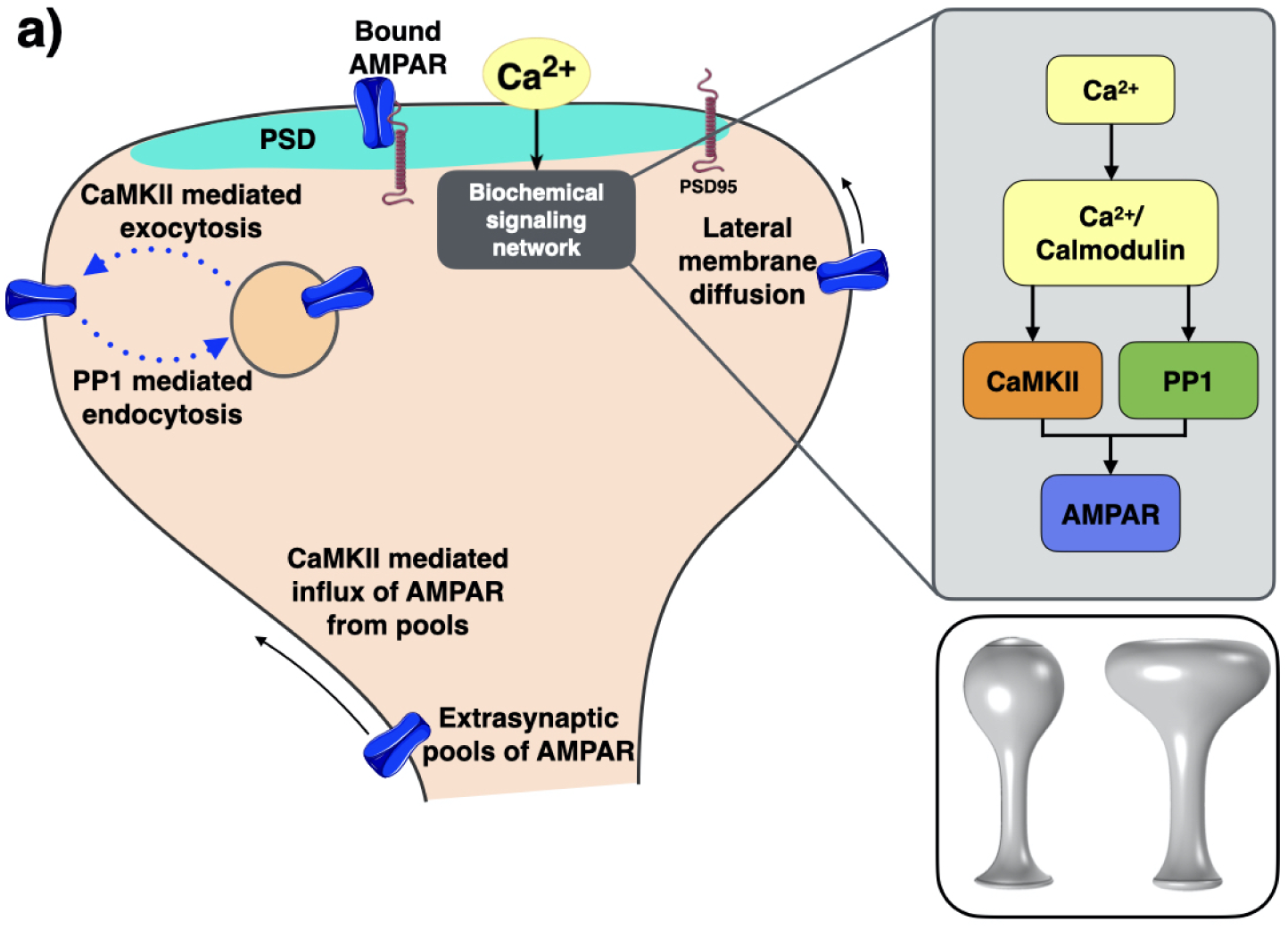
a) A combination of geometric, biochemical, and transport phenomenon factors influence AMPAR dynamics at the PSD during LTP. A biochemical network triggered by calcium influx into the spine head leads to changes in AMPAR density at the PSD. Various mechanisms govern AMPAR trafficking to the PSD, including CaMKII-mediated exocytosis, PP1-mediated endocytosis, lateral membrane diffusion, and an extrasynaptic pool of AMPAR located at the base of the spine neck. Endo/exocytosis can take place anywhere on the plasma membrane. PSD95 binds AMPAR in the PSD region to form bound AMPAR. Grey inset: Biochemical signaling involves the mutual activation of CaMKII and phosphatases that compete to influence AMPAR. Bottom inset: Idealized dendritic spines represent thin (left) and mushroom (right) spines.

### Model assumptions

- **Geometries:** We utilize both idealized and realistic geometries of thin and mushroom spines. Idealized geometries were taken from [67] and scaled to different volumes representative of average dendritic spine volumes [43, 68]. Realistic geometries were meshed from EM image stacks of a hippocampal dendritic segment [69, 70].
- **Timescales:** We focus on AMPAR dynamics on the 5-minute timescale to focus on changes during early LTP [27, 57]. Longer timescale dynamics involve other molecular pathways, including those associated with volume change during LTP, that are outside the scope of this work [57].
- **Reaction types:** The majority of the signaling reactions are either mass action or enzymatic Hill functions. Certain reactions are custom and have explicit time dependent components such as the equations for calcium influx and CaMKII-mediated AMPAR influx which have time dependent decay components. The CaMKII-mediated AMPAR influx through the spine neck base is a flux boundary condition. As a result there is no mass conservation for AMPAR in the system with influx. The model reactions are the same as those presented in [29] but translated to a spatial reaction-diffusion systems from a compartmental model. In some cases, the reactions occurring at the membrane were modeled using boundary conditions.
- **Kinetic parameters:** Reaction rates are taken from previous studies or approximated from experimental data when possible [29].
- **Boundary conditions:** The PSD is defined at the top of the spine head, and the plasma membrane is defined as all of the membrane including the PSD but excluding the bottom of the spine neck. The stimulus to the system involves a Ca^2+^ influx through the plasma membrane as a membrane boundary condition [43]. Cytosolic AMPAR can be exchanged with membrane AMPAR through a flux boundary condition on the whole plasma membrane except at the bottom of the spine neck. All other volumetric species have no flux boundary conditions on the plasma membrane. All membrane species have no flux boundary conditions at the base of the spine neck except for membrane AMPAR, which has a Dirichlet boundary condition that is dependent on both active CaMKII concentration and time. Therefore, there is no mass conservation for the AMPAR species when there is a boundary flux at the spine neck base representing influx from an extrasynaptic pool of AMPAR. We refer the reader to S1.1 and Table S6, Table S7, and Table S8 in the supplemental material for more information.

### Modular construction of biochemical signaling network

To investigate AMPAR spatiotemporal dynamics, we construct a simplified biochemical signaling network describing AMPAR dynamics on the timescale of minutes in hippocampal pyramidal CA1 neurons with deterministic reaction diffusion equations, see Fig. 1a. We highlight the key signaling components below.

- **Model stimulus and Ca^2+^ Module:** The dendritic spine is activated by a voltage depolarization at the membrane that represents an excitatory postsynaptic potential (EPSP) and a backpropagating action potential (BPAP) offset by 2 ms. This voltage spike activates NMDAR localized at the PSD, and VSCC across the whole dendritic spine membrane. Ca^2+^ is then pumped out of the spine through PMCA and NCX pumps or is bound to other signaling species. All calcium influx and efflux reactions are taken from [43].
- **Calmodulin Module:** Ca^2+^ binds calmodulin (CaM) to form a Ca^2+^/CaM complex [58]. We model this binding as a single mass action reaction, which is a simplification that has been used previously [29, 48, 58]. Free CaM can also bind to Neurogranin (Ng). Thus Ng acts as a sink for CaM in this system.
- **CaMKII and phosphatase Module:** To investigate the controversy surrounding CaMKII dynamics [27, 48], we consider two different CaMKII and PP1 models (orange and green in Fig. 1).

**– Bistable model:** The bistable CaMKII model is designed to show bistable behavior dependent on the level of calcium influx and the relative concentrations of CaMKII and PP1 [58]. CaMKII is activated by the Ca^2+^/CaM complex, can undergo autophosphorylation, and is deactivated by active PP1. For the phosphatase cascade, calneurin (CaN) is activated by the Ca^2+^/CaM complex, and subsequently activates inhibitory-1 (I1), and I1 then activates protein phosphatase 1 (PP1) which can autoactivate itself. All the phosphatases are deactivated by active CaMKII. For our selected calcium influx and initial conditions of CaMKII and PP1, we expect a sustained high concentration of activated CaMKII following activation for this bistable model [29].
**– Monostable model:** The monostable model is designed to show exponential decay of both CaMKII and PP1, due to their rate change being linearly dependent on their own concentration. In this way, CaMKII exhibits transient activation and always has a single steady state of zero concentration. Both CaMKII and PP1 are directly activated by Ca^2+^/CaM. We fit CaMKII and PP1 decay dynamics to both experimental and other modeling results [27, 29, 71, 72].
• **AMPAR signaling and trafficking Module:** The final module (blue in Fig. 1) captures AMPAR dynamics. Free AMPAR is modeled to be present on the entire plasma membrane and can be bound by PSD95 in the PSD region to become bound AMPAR with a slower diffusion coefficient. Thus, bound AMPAR is localized to a prescribed PSD membrane. Endocytosis and exocytosis can occur throughout the whole plasma membrane as boundary conditions that exchange membrane AMPAR and cytosolic AMPAR through CaMKII mediated exocytosis and PP1 mediated endocytosis [48], see S1.1.3, S1.2, and Table S6 and Table S7. Active CaMKII also mediates the influx of membrane AMPAR from an extrasynaptic pool modeled as a membrane flux at the base of the spine neck [22, 23], see Table S8.

The signaling networks used in this model can be found in [29] and [43], and are described in more detail in the Supplemental Material.

### Governing equations

We construct a system of partial differential equations representing the reaction-diffusion dynamics of signaling species in three different spatial compartments. The spatial regions include the cytoplasm (3D volume), plasma membrane (2D surface), and the PSD membrane (a subset of the plasma membrane, 2D surface). The reaction diffusion dynamics of each species, c_i_, is given by

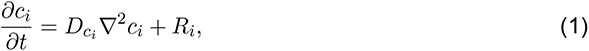

with the boundary condition given as

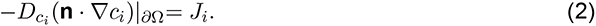

R_i_ is the reactions for species c_i_ in its respective compartment. J_i_ is the boundary condition for species c_i_. For no flux boundary conditions, J_i_ is zero. For a volumetric species, its reaction diffusion dynamics occur within the 3D volume and boundary conditions are on its encompassing 2D surface. For a membrane species, its reaction diffusion dynamics occur on its 2D surface and boundary conditions are on any encompassing 1D lines, which in this system is just the base of the spine neck. The specific equations, initial conditions, and parameters can be found in the Supplemental Material, specifically Table S4, Table S5, Table S7, Table S6,Table S1,Table S2, and Table S8. Because the spine geometry can affect the initial steady state of the system, each simulation is run to steady state without a stimulus to achieve a geometry specific steady state, see Table S3 and Table S12.

### Spine model geometry

We modeled both 2D axisymmetric idealized spines and 3D realistic spines reconstructed from EM images [69], as isolated compartments. Idealized spines were modeled after characteristic thin and mushroom spines in hippocampal pyramidal neurons, see inset in Fig. 1 and [67], and realistic spines shown in Figure 6a were selected from the dendritic segment shown in Figure S6a. A range of spine sizes were explored for both idealized and realistic geometries, with care taken to select a variety of realistic spine morphologies, see Table S10 and Table S11. The plasma membrane was modeled as the whole spine surface except for the bottom base of the spine neck, and the PSD regions were labelled at the top of the spine head for idealized geometries and as denoted during segmentation for the realistic spines. We quantify the various spine morphologies in terms of their volumes (vol) and their volume to surface area (SA) ratio.

### Parameter variations

To investigate the role of biochemical signaling, we consider two different biochemical signaling networks for CaMKII and PP1 - the bistable and monostable models described above. We also consider different trafficking knockout conditions. Specifically, we vary endocytosis and exocytosis, an active CaMKII-dependent and time-dependent extrasynaptic AMPAR influx, and membrane diffusion of free membrane AMPAR. Computationally, these trafficking knockout cases involve removing relevant terms from the model, see Table S9. We also consider stimulus frequency by using multiple active calmodulin pulses as input into the system.

### Numerical methods

All simulations were run in COMSOL Multiphysics, a commercial finite-element software. The general form and boundary partial differential equations (PDEs) interface was used to solve the governing equations. For the various simulations, a COMSOL generated mesh of “extra fine” or “extremely fine” was selected with several boundary layers added to the membrane. The absolute tolerance was lowered to 0.0001 for all simulations. Analysis and plotting were performed in MATLAB. The model files can be found on GitHub under RangamaniLabUCSD/AMPAR-PDE-Bell.

## Results

We develop two separate models of CaMKII and PP1 dynamics to explore both bistable dynamics that lead to elevated activated CaMKII concentration and monostable dynamics that lead to transient CaMKII dynamics, see Figure S1 for both CaMKII and PP1 temporal dynamics. We consider bound AMPAR dynamics at the PSD as both a receptor density and total receptor number as the readout of the model. Total receptor number is obtained by integrating receptor density over the PSD membrane area. We investigate three different trafficking mechanisms as shown in Fig. 1a - a CaMKII-mediated influx of membrane AMPAR from an extrasynaptic pool of membrane-bound AMPAR at the base of the spine neck (influx term), lateral membrane diffusion of AMPAR, and CaMKII and PP1 mediated exocytosis and endocytosis of AMPAR, respectively. We vary these trafficking conditions in five cases - 1. a control case with all trafficking conditions, 2. no influx at the spine neck base (no influx), 3. no influx and no endo/exocytosis of AMPAR (only diffusion), 4. no endocytosis/exocytosis (no EnEx), and 5. no lateral membrane diffusion of AMPAR (no diffusion).

### Bound AMPAR depends on spine volume to surface ratio

We simulated the biochemical network involving calcium influx through localized receptors and channels, Calmodulin activation, subsequent CaMKII and phosphatase activation, and AMPAR change. We saw that at these timescales, despite the localization of receptors and channels for calcium, all cytosolic species other than cytosolic AMPAR showed homogeneous spatial dynamics, Figure S1. Therefore, due to this homogeneity and the previous investigation of these signaling species [29, 43, 45, 46, 73], we will focus specifically on AMPAR spatiotemporal dynamics.

We first considered spatial and temporal plots of membrane AMPAR (Amem) for the control thin and mushroom spines to investigate how membrane dynamics of AMPAR inform bound AMPAR, Figure 2. We see that membrane AMPAR has a spatial gradient in both spine shapes and for both models because of the CaMKII-dependent influx of AMPAR at the spine neck. This gradient is apparent in both the spatial plots and in the temporal plots that highlight the Amem density difference between the PSD region and the spine neck base. The PSD region has a lower density of Amem due to its binding to PSD95 that forms bound AMPAR. We see that this gradient persists for longer in the bistable model (Figure 2a-b,e-f) compared to the monostable model (Figure 2c-d,g-h) because active CaMKII remains elevated in the bistable model which leads to a larger AMPAR influx. Similarly, the bistable model has higher maxima for Amem compared to the monostable model. Additionally, the mushroom spine, which has a larger control volume compared to the thin spine, has higher maximum Amem peaks compared to the thin spine. Thus, we see that the spatial segregation of these trafficking mechanisms, in particular the distance between the AMPAR influx at the spine neck base and PSD95 binding at the PSD, means that spine geometry can influence membrane AMPAR and thus bound AMPAR dynamics.

**Figure 2:**
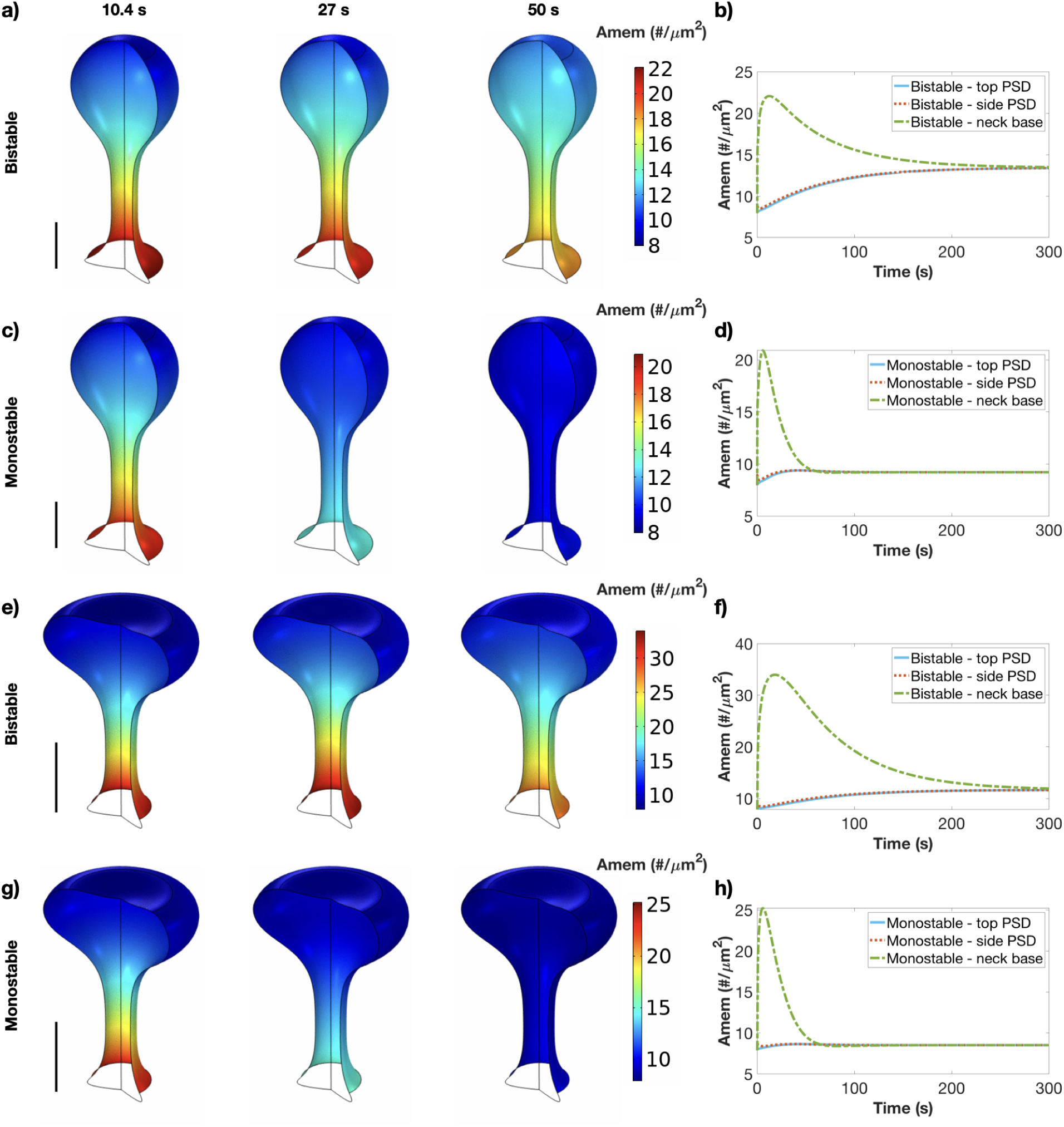
Membrane AMPAR spatiotemporal dynamics for the control thin and mushroom spines emphasize trafficking localization. a) Spatial dynamics of membrane AMPAR (Amem) in the bistable model at three time points (10.4 s, 27 s, and 50 s) for the thin control spine. Scalebar is 200 nm. b) Temporal dynamics of membrane AMPAR in the bistable model for the thin control spine at three locations (center of the PSD, side of the PSD, and base of the spine neck). c) Spatial dynamics of membrane AMPAR (Amem) in the monostable model at three time points (10.4 s, 27 s, and 50 s) for the thin control spine. Scalebar is 200 nm. d) Temporal dynamics of membrane AMPAR in the monostable model for the thin control spine at three locations (center of the PSD, side of the PSD, and base of the spine neck). e) Spatial dynamics of membrane AMPAR (Amem) in the bistable model at three time points (10.4 s, 27 s, and 50 s) for the mushroom control spine. Scalebar is 500 nm. f) Temporal dynamics of membrane AMPAR in the bistable model for the mushroom control spine at three locations (center of the PSD, side of the PSD, and base of the spine neck). g) Spatial dynamics of membrane AMPAR (Amem) in the monostable model at three time points (10.4 s, 27 s, and 50 s) for the mushroom control spine. Scalebar is 500 nm. h) Temporal dynamics of membrane AMPAR in the monostable model for the mushroom control spine at three locations (center of the PSD, side of the PSD, and base of the spine neck).

We considered an idealized thin spine and mushroom spine with volumes representative of average thin and mushroom spines, respectively. Typically thin spines tend to be smaller in volume than mushroom spines, so the control thin spine is approximately 7 times smaller than the control mushroom spine. The thin spine and mushroom spine show similar kinase, phosphatase, and bound AMPAR temporal dynamics, so we plot active CaMKII, active PP1, and bound AMPAR at the top of the PSD of the thin control spine for both the monostable and bistable models Figure 3a-c. We see that the bistable model has an elevated CaMKII concentration and transient PP1 concentration, while the monostable model has transient dynamics for both CaMKII and PP1 Figure 3a-b. This translates to a higher bound AMPAR steady state for the bistable model compared to the monostable model due to the elevated CaMKII dynamics that drive exocytosis and AMPAR influx, Figure 3c. These dynamics are extensively studied in [29].

**Figure 3:**
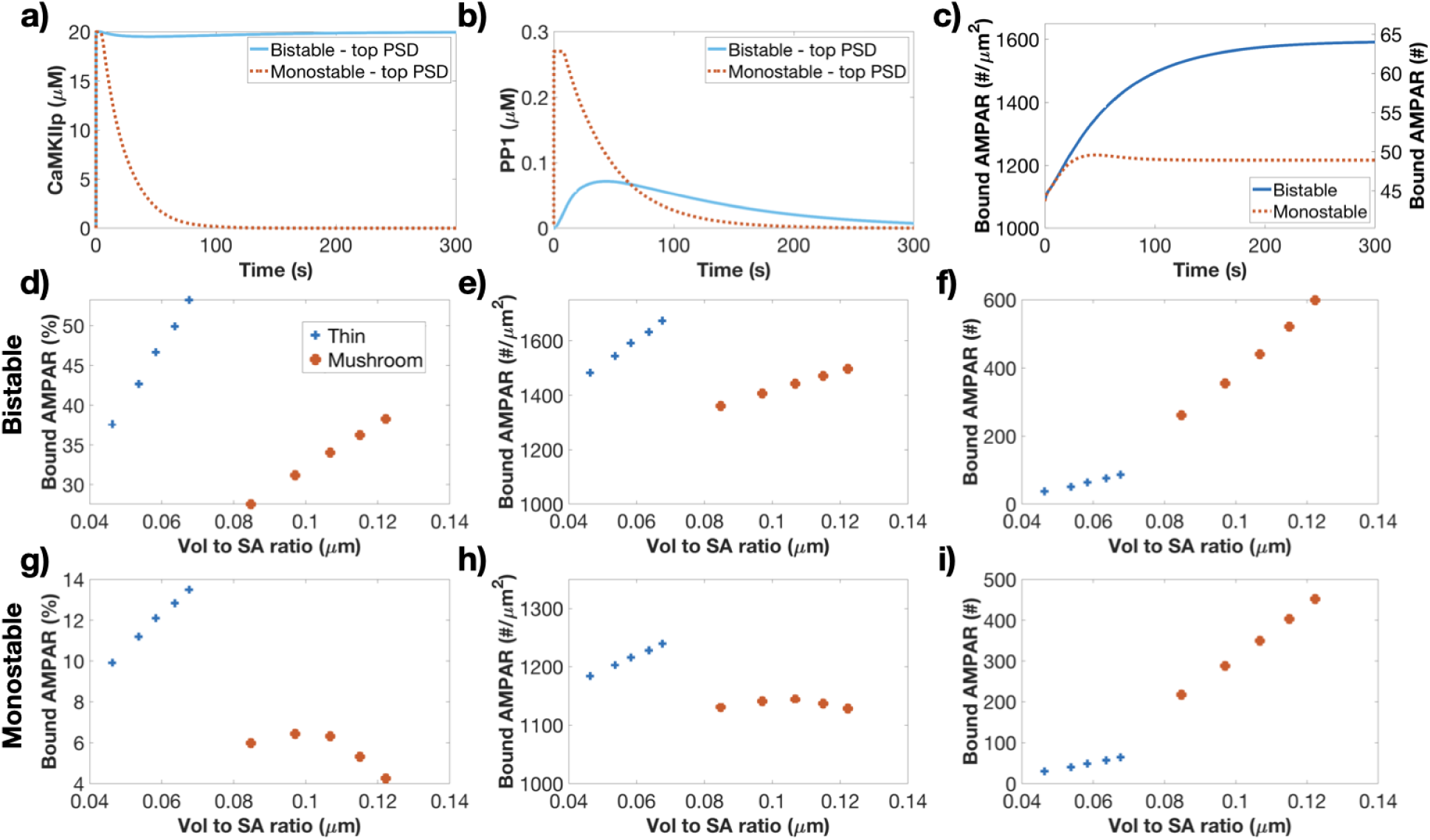
Bound AMPAR dynamics for thin and mushroom spines of different sizes show a resetting between spine shapes. Temporal dynamics of active CaMKII (a), active PP1 (b), and bound AMPAR (c) for both the bistable and monostable (blue and red lines, respectively) at the top of the PSD for a control thin spine. We consider the steady state of bound AMPAR at 300 s for each spine for both the bistable and monostable models as a percent change from steady state, receptor density, and total receptor number versus volume to SA ratio for the bistable model (d-f) and monostable model (g-i). The blue and red points denote the thin and mushroom spines, respectively.

We investigated how spine volume can influence these AMPAR dynamics by increasing and decreasing the control thin and mushroom spine volumes by 50% to explore the dendritic spine size space. We considered the bound AMPAR steady state for the different sizes of the two spine shapes for both models. As noted above, each simulation was run to steady state before applying the model stimulus, resulting in slightly different bound AMPAR steady states for each spine geometry. We plot the initial conditions for bound AMPAR for all spine geometries as a receptor density and total receptor number (Figure S4) and we see that while the thin spines tend to have higher initial receptor densities, those densities translate to much smaller total receptor numbers compared to the mushroom spines due to the smaller PSD surface areas.

We ran simulations in these various spine geometries and considered the steady state value for bound AMPAR as a percent change from steady state, receptor density, and total receptor number, and plot them against volume to surface area ratio for the different spines geometries and two different biochemical models, Figure 3d-i. For the bistable model (Figure 3d-f), plots of bound AMPAR density and percent change versus volume to surface area ratio show similar trends; as the thin spines (smaller, blue dots) and mushroom spines (larger, red dots) increase in volume to SA ratio within their shape group, the steady state bound AMPAR density (and percent change) increases, Figure 3d-e. However, when transitioning from the thin spines to the larger mushroom spines, the bound AMPAR density (or percent change) drops, resetting to a lower density (or percent change) that then increases as the mushroom spines gets larger. For the monostable model (Figure 3g-i), plots of bound AMPAR density and percent change versus volume to surface area ratio show a similar reset between the thin and mushroom spines (Figure 3g-h); however within the mushroom spines (red dots), there is a nonmonotonic change in receptor density and percent change as the volume to SA ratio increases. When considering total receptor number (Figure 3f,i), both the bistable and monostable models show similar trends with increasing receptor numbers for increasing volume to surface area ratio which is to be expected from the initial conditions. There is a discontinuity of the trend between the different spine shapes when considered versus volume to SA ratio. We see that these trends hold when we plot the same bound AMPAR readouts versus volume, Figure S3.

For completeness, we plot the temporal dynamics of bound AMPAR for each spine shape and size for the two different models (Figure S3). We see that all spine sizes show similar monotonic increasing trends for the bistable model, while all spine sizes show a transient peak and subsequent reduction to a lower steady state for the monostable model.

Therefore, while the bound AMPAR temporal dynamics all show similar trends within model type, bound AMPAR steady state shows shape, volume, and model-dependent trends. We found that there is a resetting of the bound AMPAR density (or percent change) between the two spine shapes, despite the increase in spine volume (and vol to SA ratio) between the thin and mushroom spines. This trend matches the pattern found for synaptic weight updates based on Ca^2+^ influx into dendritic spines of different sizes and geometries [46]. Therefore, we predict that thin spines are able to increase the density of bound AMPAR density more rapidly for increases in their volume; while in comparison, mushroom spines have a slower increase in density as volume increases. This supports previous reports that thin spines are more adaptive compared to mushroom spines [36, 37]. This resetting of bound AMPAR for larger mushroom spines can act to optimize biochemical resources, preventing excessive bound AMPAR density increases for larger spines which would translate to much higher total receptor numbers. When considering total receptor number, larger spines had larger total numbers of receptors after stimulation, but also start with higher receptor levels (Figure S4). The biochemical models also show important differences with the bistable model leading to much larger percent increases in bound AMPAR compared to the monostable model (*∼*55% increase vs *∼*14% increase, respectively). Additionally, the mushroom spines with the monostable model showed the only dynamics that were not a monotonic increase in bound AMPAR density or percent change (Figure 3g-h). It is possible that as the mushroom spines are larger in volume, there was a trade off between how quickly AMPAR could flood into the spine due to the CaMKII-dependent influx and subsequently diffuse to the PSD versus how far the AMPAR must travel and how quickly CaMKII deactivates in the monostable model. Thus, we predict that changes in bound AMPAR depend on upstream signaling dynamics (model type) and spine shape including spine volume and volume to SA ratio.

### The monostable model integrates calmodulin stimulus as a leaky integrator

We next considered how stimulus frequency can influence bound AMPAR readout. We applied active CaM pulses at different frequencies throughout the whole spine as the model input, Figure 4a. We note that we used a much smaller active calmodulin magnitude for these pulses compared to our model (see Figure S1b) to prevent rapid activation of CaMKII which would prevent us from seeing any temporal nuances. The bistable model shows no dependence on frequency, with all stimulus activating CaMKII to its maximum concentration and bound AMPAR showing similar dynamics, Figure S2. For the monostable model, CaMKII acts as a leaky integrator for all frequencies and this integration behavior translates to stepping bound AMPAR dynamics for the 0.1 Hz and 0.05 Hz stimulus, Figure 4b-f. Free AMPAR (Amem) show location dependent behavior as the influx at the spine neck base is dependent on active CaMKII, Figure 4e. These findings agree with predictions from our compartmental model [29].

**Figure 4:**
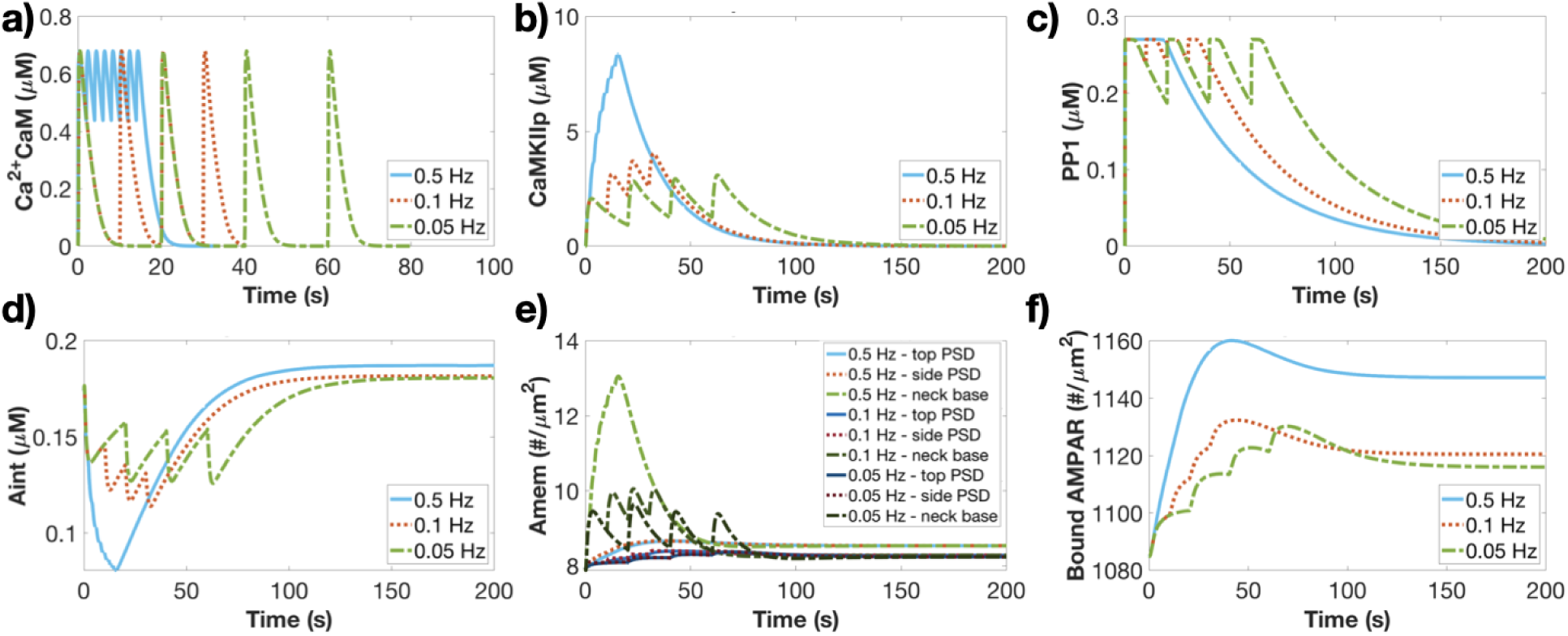
Leaky integrator temporal dynamics in the monostable model for Ca^2+^CaM pulses of different frequencies. a) Ca^2+^CaM temporal dynamics at three different frequencies (0.5, 0.1, and 0.05 Hz) serve as the input for the monostable model. Temporal dynamics of active CaMKII (b), active PP1 (c), cytosolic AMPAR (d), membrane AMPAR (e), and bound AMPAR (f) for three different frequencies plotted at the top of the PSD for the monostable model. Membrane AMPAR (e) is plotted at the top of the PSD, side of the PSD, and at the neck base.

### Trafficking knockout cases mimic trends in the control trafficking cases

We next investigated the contributions of different trafficking mechanisms by systematically knocking out various terms. What role does each trafficking term play in bound AMPAR temporal and steady state behavior? And how does spine geometry influence those trafficking roles? Specifically, we consider five different cases - 1. a control case with all trafficking conditions (all), 2. a case with no influx at the spine neck base (no influx), 3. a case with no influx and no endo/exocytosis of AMPAR (only membrane diffusion), 4. a case with no endocytosis/exocytosis (no EnEx), and 5. a case with no lateral membrane diffusion of AMPAR (no membrane diffusion). We consider the temporal dynamics of both the thin and mushroom control spines for the bistable and monostable model for these various trafficking variations, Figure 5a-d. Both spine shapes show similar bound AMPAR temporal dynamics within each biochemical model.

**Figure 5:**
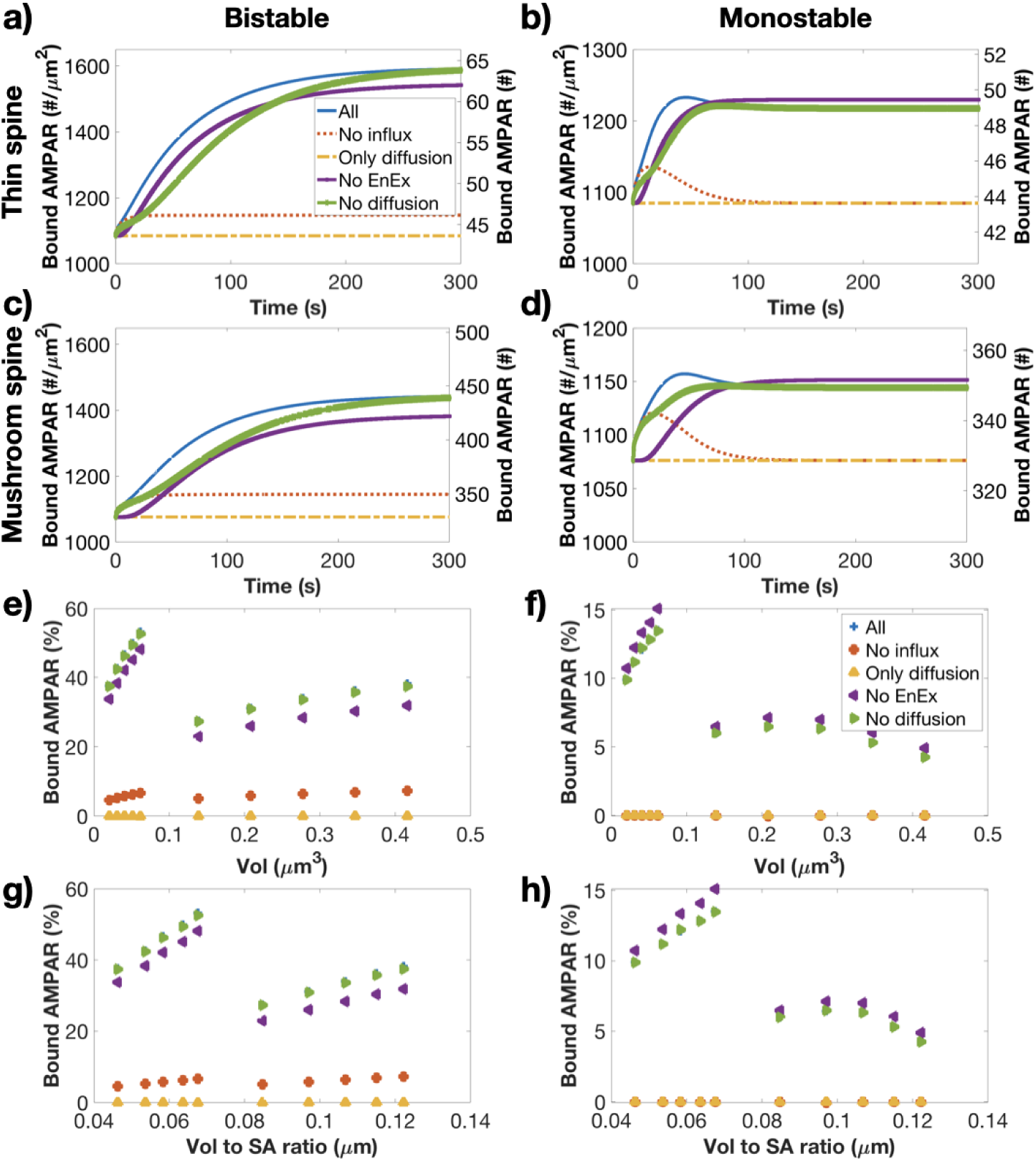
Bound AMPAR dynamics and steady state behavior for different trafficking knockout conditions. Temporal dynamics of bound AMPAR at the top of the PSD for the thin control spine for different trafficking knockout cases for the bistable (a) and monostable (b) models. Temporal dynamics of bound AMPAR at the top of the PSD for the mushroom control spine for different trafficking knockout cases for the bistable (c) and monostable (d) models. Trafficking knockout legend located in a for panels a-d. Percent change in bound AMPAR steady state from initial condition for all spine volumes for the different trafficking knockout cases for the bistable (e) and monostable (f) models. Percent change in bound AMPAR steady state from initial condition for all spine volume to SA ratios for the different trafficking knockout cases for the bistable (g) and monostable (h) models. Trafficking knockout legend located in f for panels e-h.

We will review the bistable model behavior first, Figure 5a,c. Bound AMPAR in the all trafficking (all, blue) condition increases quickly and plateaus gradually as the CaMKII-mediated influx continues since CaMKII remains activated in the bistable model for these initial conditions. When there is no influx (red), we see a similar rapid increase like the control case but then a quick plateau to a lower steady state value. In the case with only membrane diffusion (yellow), there is no change in bound AMPAR, verifying that the system begins at steady state. In the absence of endo/exocytosis (purple), we see a clear temporal delay in bound AMPAR increase before it plateaus in the same manner as the control case but at a slightly reduced steady state. In the absence of membrane diffusion (green), we see a fast initial increase but then slower increase dynamics before the system eventually reaches the same steady state as the control case.

The monostable model leads to different behavior compared to the bistable model, Figure 5b,d. The control case (all, blue) rapidly increases to a peak, slightly decreases and then plateaus to a steady state that is significantly lower than the bistable model. This is due to the transient CaMKII dynamics which means that there is a significantly smaller influx of extrasynaptic AMPAR at the spine neck for the monostable model. The no influx case transiently increases and then returns to its initial steady state (red). This supports the concept that bound AMPAR requires a consistent forcing term (sustained elevated CaMKII to force exocytosis) or a change in available AMPAR to reach a new steady state [29]. Similar to the bistable model, the case with only membrane diffusion (yellow) shows no change from initial condition. The case without endocytosis/exocytosis (purple) shows a delayed increase in bound AMPAR but smoothly reaches a new steady state value that exceeds the steady state for the control case. In the absence of membrane diffusion (green), similar to the bistable model, we get an initial increase like the control case but then a slower gradual increase to the same level as the control case.

Interestingly, the case without endocytosis/exocytosis in the monostable model has a higher steady state than the control case. In the control case, the monostable model has an increase in internal AMPAR following stimulus, with Aint steady state increasing from its initial condition (compared to the bistable model decreasing from its initial condition). Without the ability to undergo endocytosis, the system has more available membrane AMPAR to convert to bound AMPAR, leading to the elevated steady state compared to the control case.

We quantified these time dynamics by considering the time to half max for each trafficking case in the thin and mushroom control spines for both model types, Table 1. We see that the monostable model consistently has faster time to half max compared to the bistable model, for both spine shapes. Considering the different knockout cases, we conclude that the initial rapid increase in bound AMPAR is due to endocytosis and exocytosis. We also predict that the rate of increase of bound AMPAR after approximately 25 seconds is membrane diffusion dependent and therefore modifications to AMPAR membrane diffusion could modulate the later stage dynamics of AMPAR increase. Additionally, the steady state of bound AMPAR depends on the total amount of AMPAR available to the system, including membrane AMPAR from the influx and cytosolic AMPAR. Assuming available endocytosis and exocytosis locations, even without membrane diffusion, bound AMPAR can increase to the same steady state as the control case.

**Table 1:** Time to half max [s] for different trafficking cases in thin and mushroom control spines

|  | **No influx** | **Only diffusion** | **No En/Ex** | **No diffusion** | **Control** |
| --- | --- | --- | --- | --- | --- |
| Thin bistable | 2.7 | 0 | 52.6 | 76.8 | 43.6 |
| Thin monostable | 1.8 | 0 | 22.7 | 25.5 | 11 |
| Mushroom bistable | 6.1 | 0 | 74.8 | 81 | 51.5 |
| Mushroom monostable | 2.9 | 0 | 41.3 | 9.3 | 9.0 |

Values for the control volume spines. The thin control spine volume is $0.040655 \mu\text{m}^3$ and the mushroom control spine volume is $0.27738 \mu\text{m}^3$ . All times are in seconds.

Next, we investigated whether spine size can affect these trafficking trends. We consider the percent change in bound AMPAR steady state versus both volume and vol to SA ratio for spines of different sizes for all trafficking cases, Figure 5e-h. We found that the patterns versus vol and vol to SA ratio observed previously for the control case hold for the various knockout cases. In particular, the no influx case shows no change from initial density for the monostable model but marginal increases in the bistable model (red dots). The only diffusion case shows no change in bound AMPAR for both models and all geometries (yellow dots). The no endo/exocytosis case mimicks trends with respect to geometry found in the control case, but with a slight decrease in the bistable model and slight increase in the monostable case (purple dots). The no diffusion case has the same changes in bound AMPAR steady states as the control case (green dots that overlap the blue dots). Therefore, we found that we must consider both temporal and steady state dynamics of bound AMPAR to fully understand the consequences of trafficking mechanisms.

### Realistic spines show a similar trend as idealized spines for the bistable model

We next considered if these geometric trends held for realistic spines which have much more variability and complexity to their morphologies [74]. We used spine geometries reconstructed from EM images [69] and meshed to a sufficient quality to run simulations [70]. However, due to the complexity of the geometries and thus numerical complexity, we considered a simplified model setup. Specifically, due to the spatial homogeneity of upstream signaling species, we only modeled the AMPAR species and PSD95 with a temporal input of active CaMKII and PP1 from the thin control spine simulation results. We also localized membrane endocytosis and exocytosis to the PSD membrane, which delays the rise to steady state but should achieve similar steady state as if endocytosis and exocytosis existed everywhere. We considered the bistable model in these realistic spine geometries because the idealized bistable model spines showed fairly large percent increases and clear trends, Figure 3. We picked seven reconstructed spines of different volumes from a single dendritic segment; spines shown in Figure 6a and segment shown in Figure S6a. The different spines have a variety of morphologies with different volumes, surface areas, and PSD surface areas, Figure 6b. The simulations were run to an initial steady state, and with the CaMKII and PP1 stimulus all show increasing bound AMPAR dynamics, Figure S6b.

**Figure 6:**
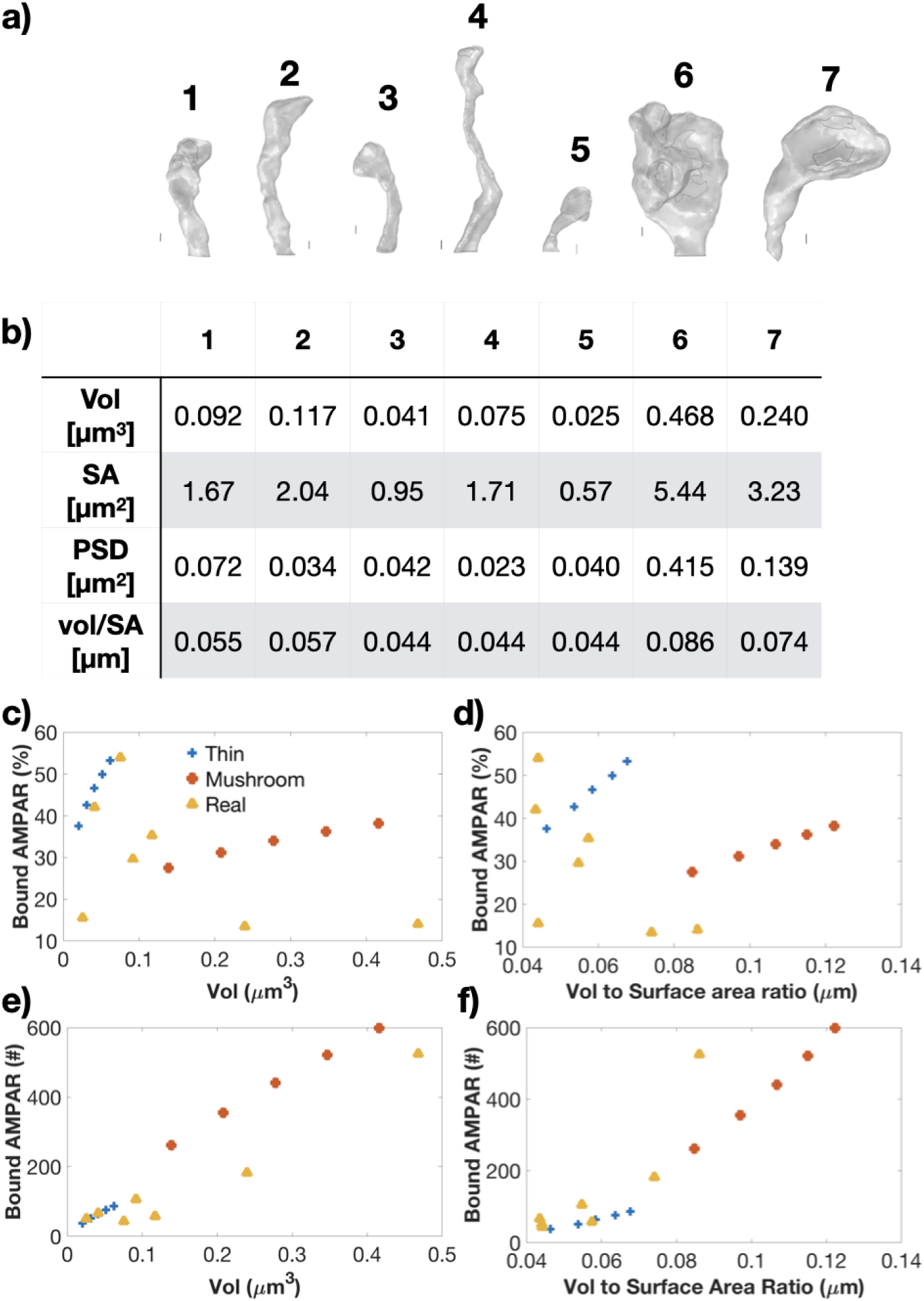
Bound AMPAR dynamics and steady state behavior for the bistable model in realistic geometries. a) Realistic dendritic spines selected from the dendritic segment reconstructed from EM data stacks in [69]. We conduct simulations for the bistable model in all the numbered spines. Scale bars next to each individual spine correspond to 100 nm. PSDs are denoted by light grey lines on the individual spines. b) Geometric parameters including volume, surface area, PSD surface area, and volume to surface area ratio for the various realistic spines. Percent change in bound AMPAR steady state for idealized and realistic spines versus spine volume (c) and volume to SA ratio (d). The thin spines are in blue, mushroom spines in red, and realistic spines are in yellow; legend located in c for panels c-f. Steady state of total number of bound AMPAR at the PSD for idealized and realistic spines versus spine volume (e) and volume to SA ratio (f).

We compared the percent increase in bound AMPAR steady state in the realistic spines to the ideal thin and mushroom spines, and observe a noisier trend, Figure 6c-d. Specifically, when plotted against volume or volume to SA ratio, in the smaller vol (or vol to SA) regime, there is a faster increase in percent increase as the spines increase in volume (or vol/SA). The percent increase then resets around a particular value for volume or vol/SA, before increasing again for larger volumes (or vol/SA). Therefore, it appears that realistic spines follow a similar trend to the idealized spines; within a smaller volume (or vol to SA ratio) regime (below 0.1 µm^3^ or 0.07 µm), percent increase in bound AMPAR increases quickly as the geometric parameter increases, then there is some resetting regime where the increase change falls, and then the percent change increases with increasing geometric parameter again but at a slower rate. However the realistic spines show more variability in the model outcomes because of their shapes. For example, comparing realistic spines 3, 4, and 5, these spines have different morphologies and volumes, but almost exactly the same volume to surface area ratio. Therefore, the geometric parameter we consider will show a different trend for bound AMPAR increase Figure 6c-d.

We last considered the total receptor number by integrating receptor density across PSD surface area for the realistic spines and idealized spines versus volume and volume to SA ratio, Figure 6e-f. We see that similar to the idealized spines, the realistic spines show an increasing trend in total receptor number versus both volume and volume to SA ratio.

## Discussion

AMPAR dynamics are vital for synaptic plasticity, synaptic transmission, and proper neuronal function. The importance of AMPAR in LTP is well established and much work has been done to investigate the signaling pathways and trafficking modalities related to AMPAR behavior [7, 27, 48, 50, 54]. However, how all these various factors interact to modulate AMPAR dynamics remains unknown. In this work, we proposed that AMPAR dynamics depend on a combination of trafficking modalities and a complex interplay between spine morphology and signaling networks. We investigated these dynamics by considering two different biochemical models for the underlying kinase and phosphatase interactions (bistable and monostable) in idealized spines of two different shapes (thin and mushroom). We systematically varied the volumes of these two idealized shapes to explore the geometric parameter regime (Figure 2 and Figure 3) and varied the biochemical stimulus frequencies to consider biochemical filtering (Figure 4). We then performed knockout simulations of the different trafficking mechanisms in the idealized spines of different volumes, Figure 5. Finally, we compared our predictions for the bistable model in idealized spines with realistic spine morphologies, Figure 6. We make several insights based on these simulations which we will discuss below.

The first insight is that that upstream cytosolic signaling species show homogeneous spatial dynamics at these extended timescales (Figure S1) and mimic the dynamics seen in our previous compartmental model results [29]. However, we did see spatial heterogeneity in cytosolic AMPAR and the membrane bound species. For the AMPAR species, we found that the influx of membrane AMPAR at the spine neck leads to a transient elevation in cytosolic and membrane AMPAR at that location, but given enough time, both species return to a single steady state, Figure S1h-i and Figure 2. However, if more specific localization of cytosolic species is considered, the role of cytosolic nanodomains might come into play [28].

The second insight is that, similar to the previous compartmental model results, the monostable model exhibits leaky integrator dynamics in a full spatial system while the bistable model did not, see Figure 4 and Figure S2; compare to compartmental model in [29]. In this spatial system, there are more distinct integration steps with the lower frequency pulses compared to ODE model, which could be attributed to diffusive effects as the system has to equilibrate across the whole spine geometry. The time needed to spatially equilibrate can also been seen in the overlap between the 0.1 Hz and 0.05 Hz cases, where the the lower frequency cases causes a higher delayed peak of bound AMPAR that is comparable to the 0.1 Hz case, but then drops to lower steady state. Both these first two insights show the importance of spatial modeling where reaction-diffusion coupling and localization of membrane fluxes and boundary conditions can alter signaling dynamics.

The third insight is that idealized and realistic spines showed clear trends in regards to geometric parameters for both models, with a reset regime where percent increase drops, Figure 3. Both the bistable and monostable models displayed this resetting dynamic, and the trend held in both idealized and realistic spines. There were however model dependent trends with receptor density and percent change within idealized spine shape, volume, and vol to SA ratio. The bistable model showed increasing percent change within idealized shape group as the spine size got larger; however the monostable model showed an increasing trend for the small thin spines but a nonmonotonic trend for the larger mushroom spines. Additionally, the percent changes in the bistable model are much larger than the percent changes in monostable model, which is consistent with previous studies of the system [29]. The realistic morphologies emphasized the resetting between smaller and larger spines but did not show as clear trends within those volume regimes, Figure 6. Therefore, the bound AMPAR response can be thought of as three geometric regimes; the first regime at small volumes (or vol to SA ratio) shows large percent increases in bound AMPAR, the intermediate regime shows a suppressed response, and the large vol (vol to SA ratio) regime shows increases but at a slower rate. It is still unclear why these resetting behavior occurs at these particular volume or vol to SA ratio regime, and further investigation is needed to pinpoint the significance of those regimes.

These complex realistic spines also demonstrate how spine morphologies have various geometric parameters that can modulate their dynamics and it is still unclear what parameters matter and how they affect the system dynamics. It is important to note that we consider percent increase in bound AMPAR because bound AMPAR can be read as a density or in terms of total receptors at the PSD. It is highly likely that it is important to consider both readouts since density could capture the local environment that the spine experiences while total receptor number can capture the importance of the synapse to the overall neural system. This is particularly important because small thin spines showed a larger biochemical increase in bound AMPAR and therefore an ability to dynamically adapt; while large mushroom spines showed a smaller percent increase in bound AMPAR but represent more receptors at their PSDs, which shows their biochemical stability. Therefore, we hypothesize that dendritic spines utilize different geometric parameters to modulate their responses during synaptic plasticity. In particular, we hypothesize that the division between smaller and larger spines serves to optimize resources where the geometric features of larger spines lead them to be biochemically conservative or stable compared to smaller spines. Additionally, it might be advantageous to prevent excessive bound AMPAR increase in some intermediate spine volume regime to prevent too many medium sized spines from utilizing excessive resources [75]. This supports the idea of resource optimization amongst spine clusters, where important large spines must have enough resources to uphold the neural network architecture and fundamental communication, while smaller spines need resources to form new connections; which leads to medium spines restricting their synaptic response [76, 77]. It is highly likely that additional geometric features, such as PSD area, PSD number, spine neck geometry, and internal ultrastructure such as the presence of a spine apparatus could also serve to tune the synaptic response, as has been previously hypothesized [42]. Therefore, the control of these morphological features effectively tunes the response of a spine during structural plasticity, highlighting the importance of structure-function feedback.

Finally, the fourth insight is that different trafficking modalities impact different aspects of the bound AMPAR response. In particular, breaking the bound AMPAR response at the PSD down into three temporal regimes, the fast timescale dynamics (0-20s) appear to be due to endocytosis and exocytosis, the intermediate timescale dynamics (20-200s) appear to be due to diffusion, and finally the late timescale steady state dynamics are governed by the availability of total AMPAR (cytosolic, membrane bound, bound to PSD95, and influx from extrasynaptic pools), Figure 5. Similar to previous findings [29], the system needs either a constant forcing term (such as elevated CaMKII or PP1) or AMPAR influx to change steady state bound AMPAR. There various trafficking trends held for different spine geometries.

Therefore, we conclude that bound AMPAR steady state depends on a combination of upstream model type, spine geometry including shape and size, and trafficking mechanisms. While we have been able to couple biochemical signaling, trafficking, and spine morphology, there are limitations to this model and thus many possible extensions to this work. Below, we discuss some next steps based on key findings in our work.

We find that endo/exocytosis of AMPAR plays a key role in short timescale AMPAR dynamics; however, we do not account for the timescales of the physical mechanisms of endosome fusion or fission, or the physical obstacles that could impede endosome movement within the spine [78]. AMPAR is also known to form distinct clusters with a variety of different proteins and scaffolding molecules [11, 18, 79], and the dynamics of cluster formation and stability remain debated [16, 80]. Additionally, the presence of an extrasynaptic pool, which was modeled as a source at the base of the spine neck, complicated the system as there was no longer mass balance for AMPAR. With an extrasynaptic pool, the bistable models took much longer to reach steady state than the monostable models, since active CaMKII remained elevated in the bistable model versus transient in the monostable model. Therefore, it is important to consider the role of the dendritic spine within a spine cluster along a dendrite [76, 77].

Although we have focused specifically on AMPAR dynamics in excitatory synapses of dendritic spines, we believe that this model combination of signaling dynamics and realistic cellular geometries along with knockout trafficking cases is a powerful tool to explore biochemical and morphological coupling in general. Well-mixed models and compartmental models can provide great insight into the signaling dynamics within dendritic spines [7, 58], but cannot address the inherently spatial questions associated with protein trafficking and spine geometry. As imaging techniques, image segmentation, and meshing technologies advance [70], the ability to computationally model realistic cellular geometries improves, providing more insight into the consequences of complex real cellular structure and ultrastructure. Our results highlight the need to consider more biological complexity in terms of spatial modeling both by considering realistic spine morphologies [74, 81] and by considering the localization of proteins and microenvironments [44]. We hypothesize that AMPAR dynamics depend on a combination of trafficking mechanisms to provide robust responses to synaptic stimuli. However, more complex models will be needed to investigate this hypothesis and test when these various trafficking mechanisms could fail. In addition, future work is needed to consider a variety of other biochemical signaling interactions, particularly in microenvironments like the PSD, and other biophysical interactions, such as crowding, confinement, and liquid liquid phase separation, that could influence AMPAR dynamics during LTP.

## Acknowledgements

We thank members of the Rangamani Lab for their comments and support, and specifically thank Dr. Sage Malingen and Dr. Mayte Bonilla Quintana for their edits and comments. This work was supported by a National Defense Science and Engineering Graduate (NDSEG) Fellowship to M.K.B., a Hartwell Foundation Postdoctoral Fellowship to C.T.L., and Air Force Office of Scientific Research FA9550-18-1-0051 to P.R..

## S1 Spatial model - Reaction Diffusion equations

We construct a spatial model of signaling molecule dynamics both on membrane and in the volume, where the spatiotemporal dynamics of a species C are given by the reaction-diffusion equation,

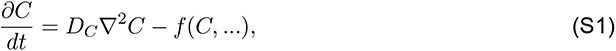

and boundary conditions are given by membrane flux in the form of

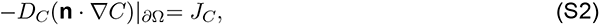

where *J_C_* captures any membrane flux reaction terms for the species C through specified boundary *∂*Ω.

### S1.1 Model Development for simplified AMPAR signaling network

The network of interest spans from calcium influx due to membrane voltage depolarization, kinase and phosphatase activation, and finally AMPAR trafficking, see Tables S1 and S2 for the initial conditions and diffusion rates of the various species. The calcium module is based on [43], the bistable model is based on [48, 58], the monostable model is inspired by experimental observations [28, 82], and the AMPAR modules are constructed based on a collection of experimental and computational observations, described in more detail below. We break the signaling network into three modules.

#### S1.1.1 Calcium influx Module

We take calcium dynamics from [43] with the exception of the cytosolic and membrane bound buffers and SERCA pumps. Calcium influx is due to activation of NMDAR localized to the PSD region and voltage sensitive calcium channels (VSCC) distributed across the whole plasma membrane. Calcium efflux is due to PMCA and NCX pumps across the plasma membrane.

#### S1.1.2 Kinase and Phosphatase Module

Calcium influx activates calmodulin which then activates CaMKII, and a phosphatase cascade. Calmodulin can also bind neurogranin. We include two different models of CaMKII and phosphatase cascade - termed bistable and monostable to differentiate. In the bistable model, CaMKII is activated by calmodulin, autophosphorylates, and is dephosphorylated by active PP1. The phosphatase cascade in the bistable model starts with calmodulin activating calcineurin which activates phosphatases I1 and PP1 in a cascade. All phosphatases are deactivated by active CaMKII, and PP1 also autoactivates itself. This model structure allows CaMKII to be perpetually active depending on the magnitude of calcium influx and initial conditions for CaMKII and PP1 [58, 83]. All reactions and reaction rates are found in Table S4.

In the monostable model, CaMKII and PP1 are both directly activated by calmodulin and their rate of decay depends linearly on their own active concentration. This model structure leads to exponential decay dynamics and a zero steady state for both CaMKII and PP1. All reactions and reaction rates are found in Table S5.

#### S1.1.3 AMPAR Module

The level of activated CaMKII and PP1 determines the rate of endo/exocytosis of cytosolic AMPAR (Aint) and membrane bound free AMPAR (Amem), which is modelled as a boundary condition. Free AMPAR exists on the whole plasma membrane while PSD95 and bound AMPAR (ABound) only exist within the PSD region. Free AMPAR within the PSD reversibly binds to PSD95 to form bound AMPAR which has a reduced diffusion coefficient. Free AMPAR enters onto the membrane through a boundary condition at the base of the spine neck representing a pool of extrasynaptic AMPAR on the dendrite. The rate of free AMPAR influx is dependent on the concentration of activated CaMKII that diffuses to the base of the spine neck. All reactions and reaction rates are found in Tables S6, S7, and S8.

### S1.2 Trafficking conditions

AMPAR trafficking during LTP is thought to include endo/exocytosis, lateral membrane diffusion, and extrasynaptic pools of AMPAR. We include all three of these sources with our computational model and systematically turn off these various features to get five trafficking cases. The first case is the control case that includes all trafficking modalities; the second case has no extrasynaptic pool; the third case has no influx and no endo/exocytosis; the fourth case has no endo/exocytosis, and finally the fifth case has no membrane diffusion of free AMPAR. The model changes for these cases are show in Table S9.

## S2 Spine geometries

Idealized spines are picked to be representative of thin and mushroom spines and the outlines are taken from [67] and used as 2D axisymmetric geometries. We varied the idealized spines to 50% and 150% of the respective control volume, see Table S10. Initial conditions for each idealized spine geometry are found in Table S3.

Realistic spine geometries were selected from a curated meshed geometry based on EM image stacks captured by Wu *et al.* [69]. The segmented EM stacks were curated with GAMer v2.0.7 and exported for import into COMSOL [70, 84]. It should be noted that a mesh of appropriate quality is necessary for successful import and for the numerical solver to run on the geometry. Realistic spine dimensions are shown in Table S11. Initial conditions for each realistic spine geometry are found in Table S12.

## S3 Simplified biochemical network for realistic spine simulations

For the realistic spines, we ran a simplified bistable signaling network. Specifically, due to the homogeneous dynamics of CaMKII and the phosphatases, the temporal dynamics of active CaMKII and PP1 from the thin control spine are taken at the top of the PSD and used as direct input into the realistic spines. Therefore, the model system only involves the AMPAR species and PSD95, with active CaMKII and PP1 temporal dynamics as the model stimulus. Additionally, to reduce model complexity, endocytosis and exocytosis were only modeled at the PSD region for the realistic spines. Because we are focusing on steady state dynamics, we do not believe that this simplification has major consequences on the readout.

## S4 Supplemental Figures

### Temporal dynamics show primarily homogeneous dynamics except for AMPAR species

We consider the temporal dynamics of key species for a thin spine of average volume, Figure S1. The temporal dynamics of key species show similar dynamics to our previous compartmental ODE model [29]. Ca^2+^ acts on a fast timescale to activate downstream species such as calmodulin (CaM), to produce Ca^2+^CaM. Calmodulin also activates Neurogranin (Ng) which acts as a CaM sink. Both models show similar early timescale dynamics (Figure S1a-c) but the different CaMKII and phosphatase models translate to very different medium timescale events (Figure S1d-g). In the bistable model, CaMKII remains elevated while the monostable model activates and decays exponentially. For both models, PP1 activates and decreases with the bistable model showing a delayed activation due to the temporal delay in the phosphatase cascade. AMPAR dynamics in the cytosol (Aint) and on the membrane (Amem) show spatial heterogeneity during early timescales but eventually reach a homogeneous steady state, Figure S1h-i. Both models show elevated Amem at the base of the spine neck (light and dark green lines) due to the Amem boundary condition that is dependent on active CaMKII activity. Cytosolic AMPAR initially rapidly decreases for both models but then increases for both at the neck base as membrane AMPAR is endocytosed due to its high density. PSD95 and bound AMPAR dynamics show opposite decreasing and increasing dynamics as expected, since PSD95 binds to membrane AMPAR to form bound AMPAR, Figure S1j-k. The bistable model produces a higher steady state of bound AMPAR and membrane AMPAR compared to the monostable model, but the monostable model has higher cytosolic AMPAR compared to the bistable steady state. Interestingly, the monostable model actually increases its cytosolic store of AMPAR after activation.

---

**Figure S1:**
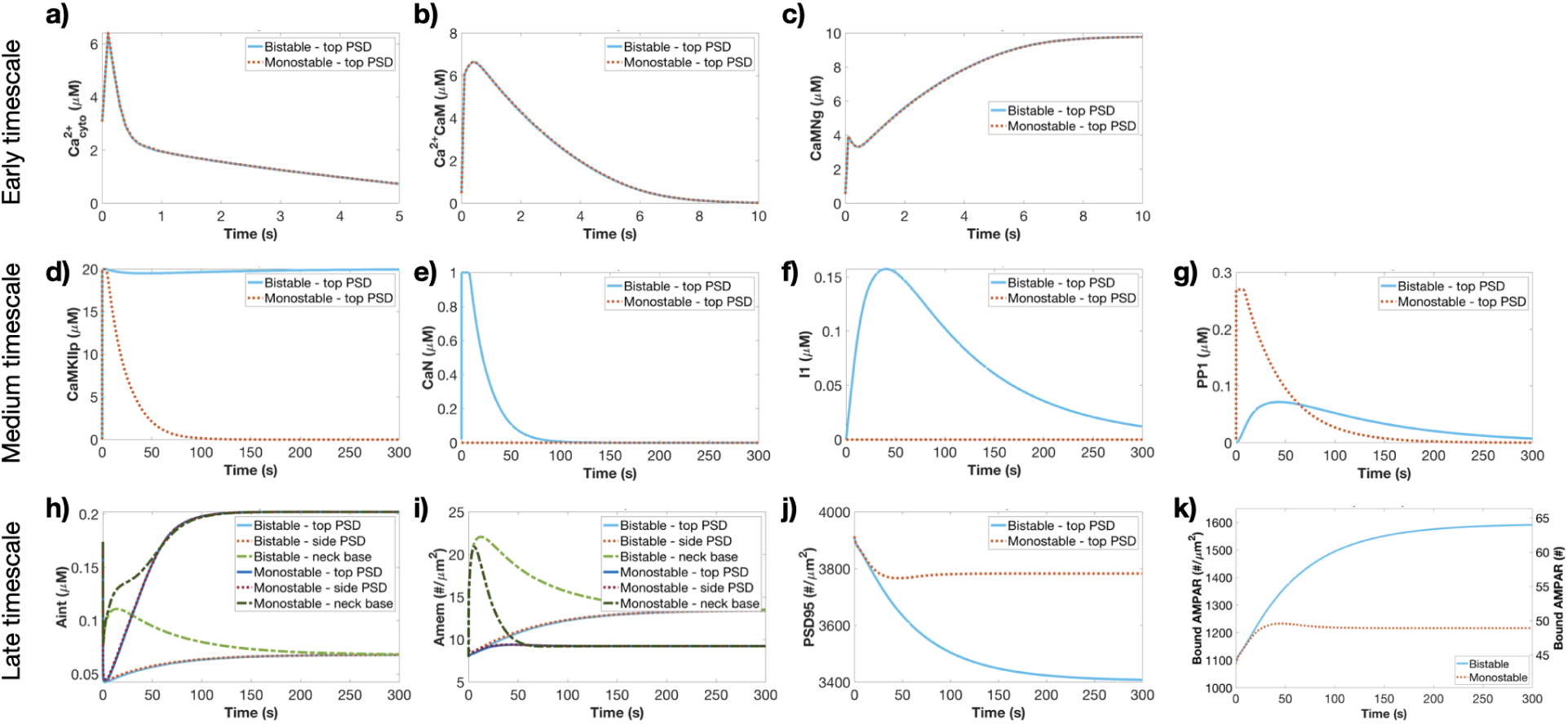
Temporal dynamics of key species in both biochemical models in a control thin spine. Temporal dynamics of the early timescale species Ca^2+^ (a), active calmodulin (Ca^2+^/CaM; b), and CaM bound Neurogranin (CaMNg; c). Temporal dynamics of medium timescale species active CaMKII (d), CaN (e), I1 (f), and PP1 (g). Temporal dynamics of late timescale species cytosolic AMPAR (Aint; h), membrane AMPAR (Amem; i), PSD95 (j), and bound AMPAR plotted as density and total receptor number (k).

**Figure S2:**
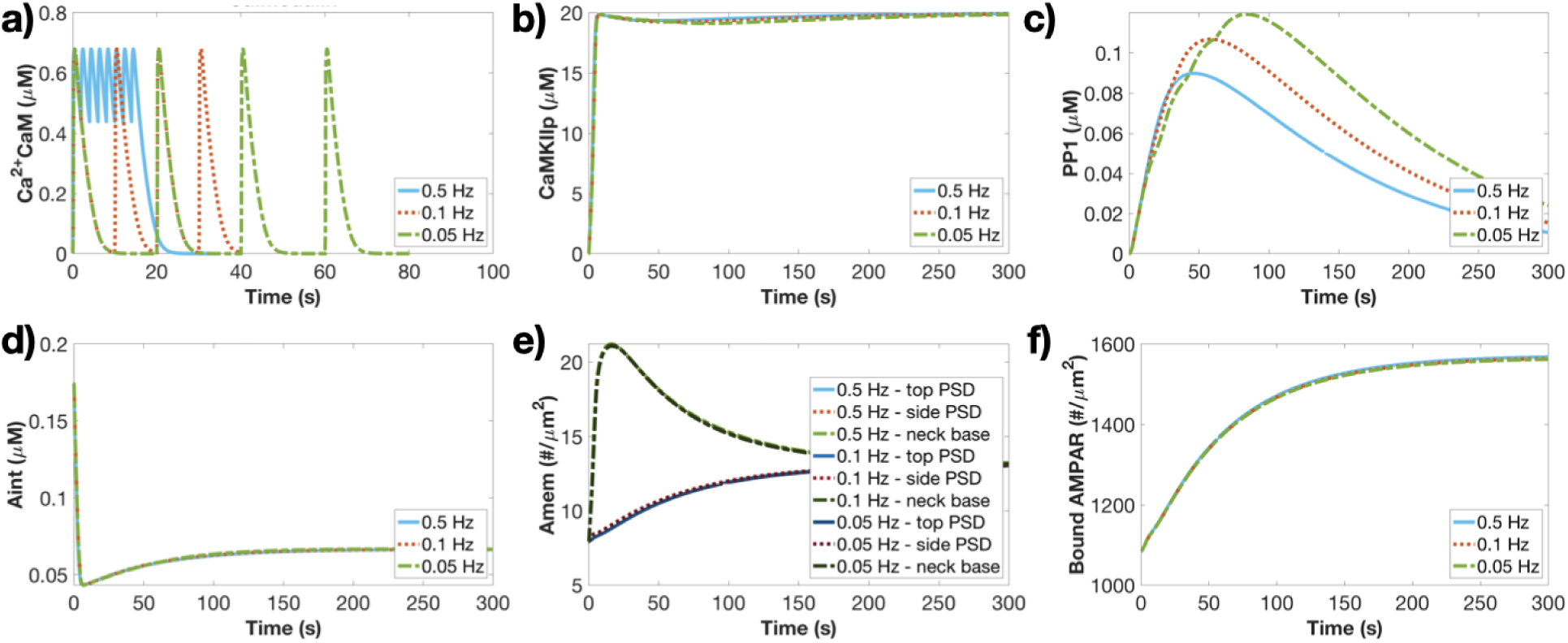
Temporal dynamics of bistable model for Ca^2+^CaM pulses of different frequencies. a) Ca^2+^CaM temporal dynamics at three different frequencies (0.5, 0.1, and 0.05 Hz) for the bistable model. Temporal dynamics of active CaMKII (b), active PP1 (c), cytosolic AMPAR (d), membrane AMPAR (e), and bound AMPAR (f) for three different frequencies at the top of the PSD of the thin control spine for the bistable model. Membrane AMPAR (e) is plotted at the top of the PSD, side of the PSD, and at the neck base.

**Figure S3:**
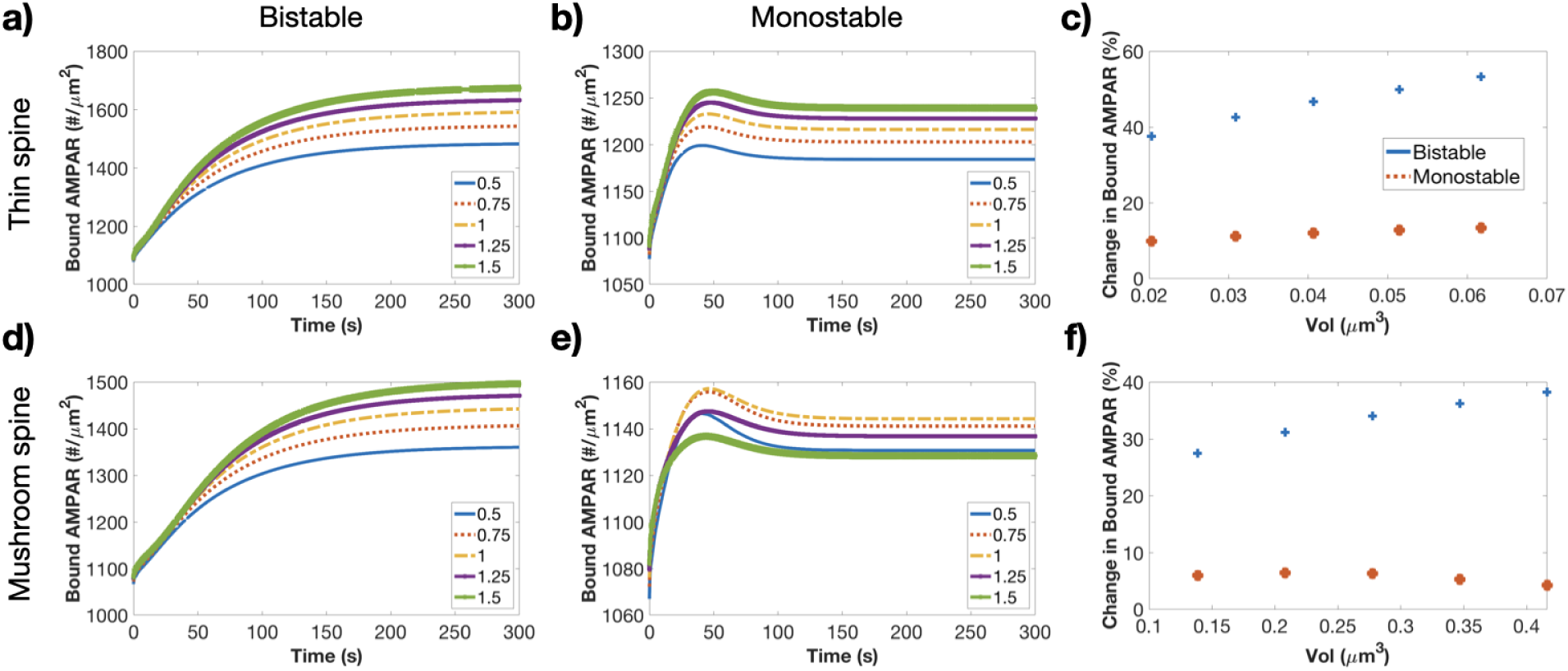
Spine volume affects bound AMPAR temporal and steady state dynamics. We plot the temporal dynamics of bound AMPAR at the top of the PSD for each size of the thin spine for the bistable (a) and monostable (b) models. c) Percent change in steady state value for each thin spine volume for the bistable and monostable models (blue and red, respectively). We plot the temporal dynamics of bound AMPAR at the top of the PSD for each size of the mushroom spine for the bistable (d) and monostable (e) models. f) Percent change in steady state value for each mushroom spine volume for the bistable and monostable models (blue and red, respectively).

**Figure S4:**
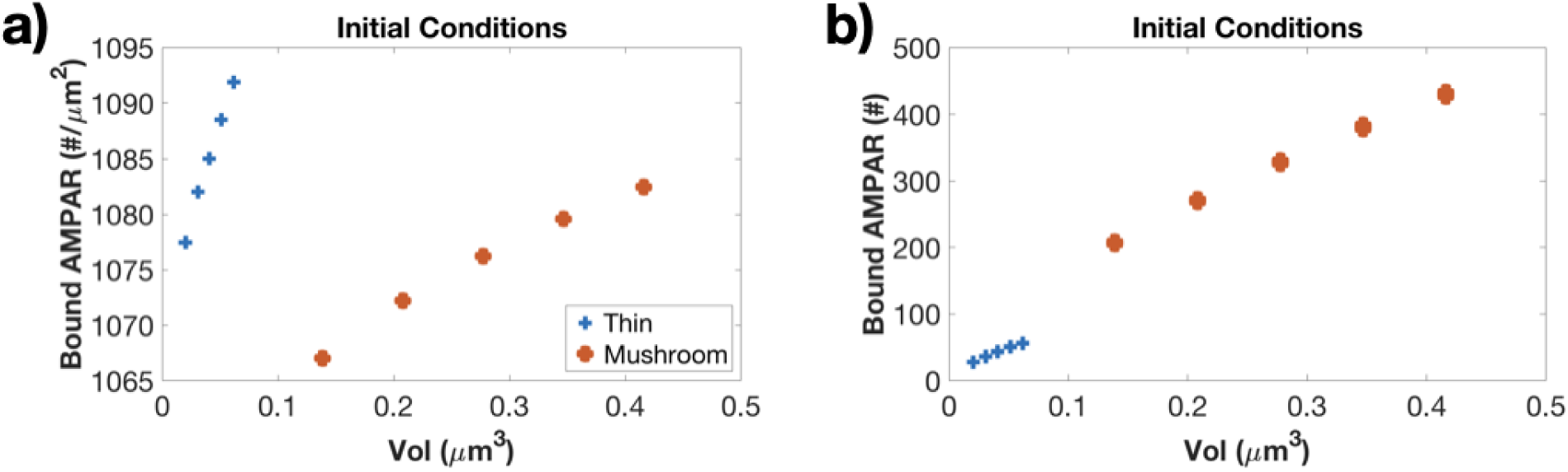
Bound AMPAR initial densities dependent on volume. Each geometry is run to its steady state without stimulus. Bound AMPAR versus volume as a receptor density (a) and total receptor number (b). Blue dots represent thin spines and red dots represent mushroom spines.

**Figure S5:**
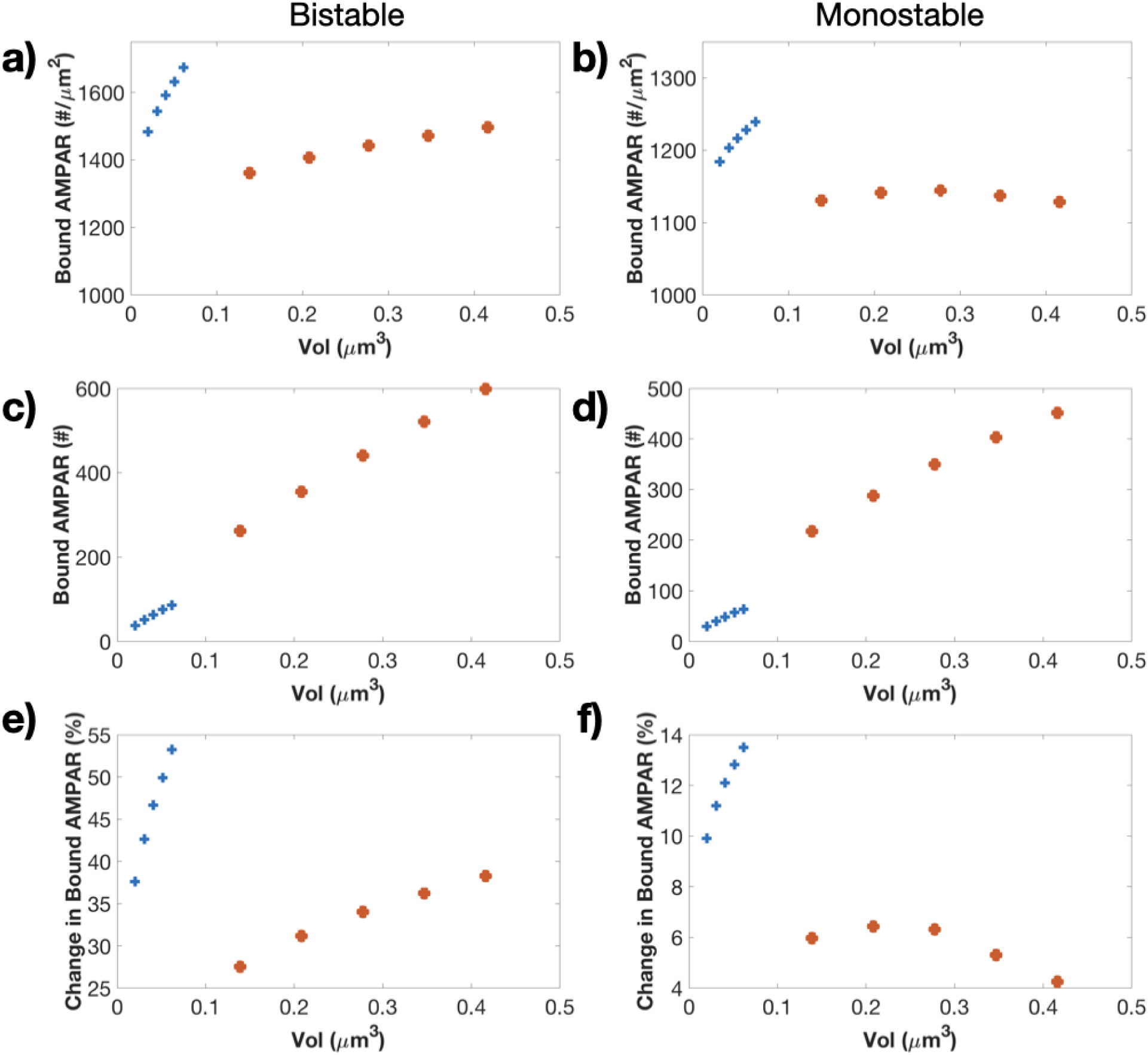
Bound AMPAR shows readout dependent trends versus volume. Bound AMPAR versus volume as a receptor density (a-b), total receptor number (c-d), and percent change in steady state value (e-f) for the bistable model (left column) and monostable model (right column).

**Figure S6:**
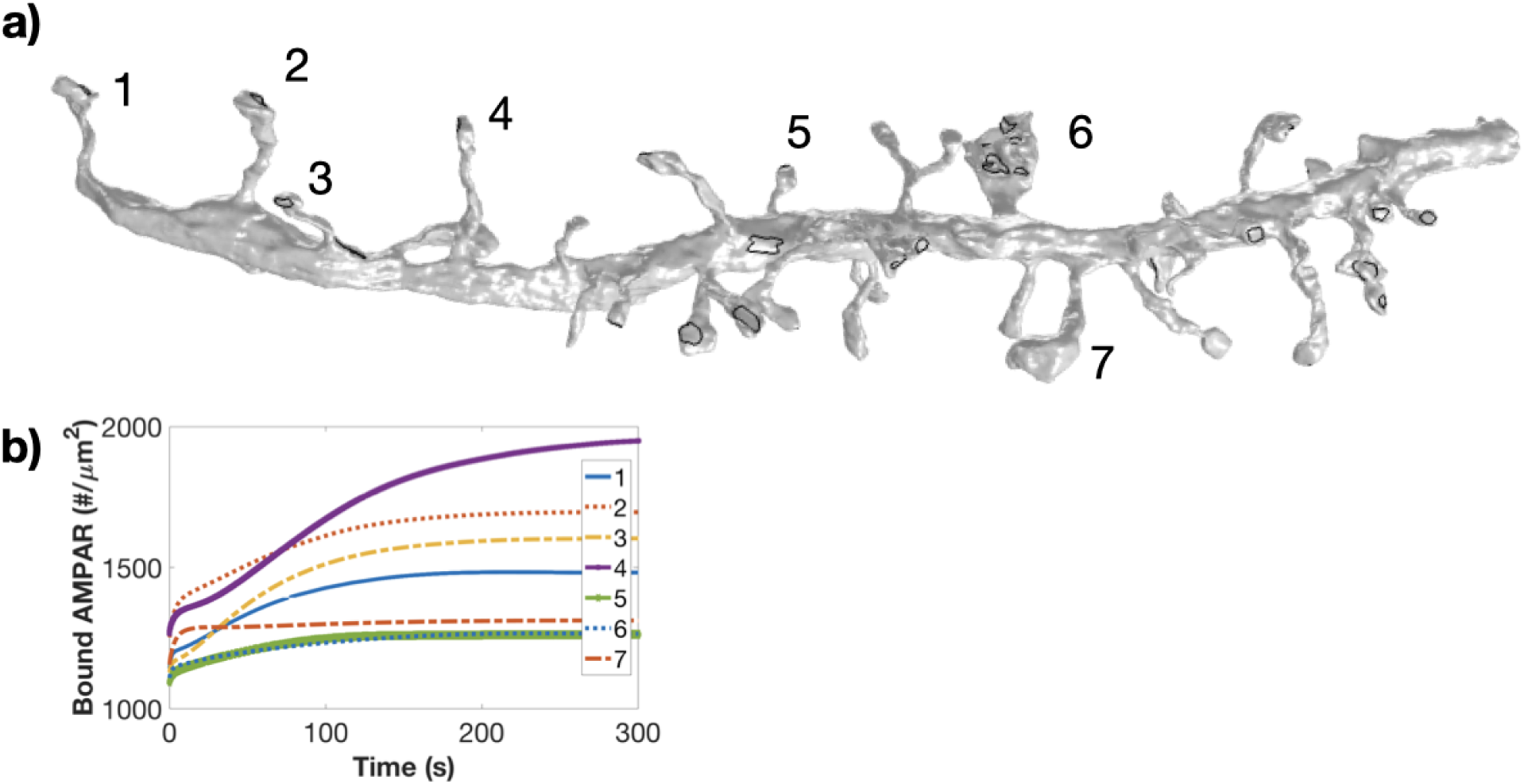
Reconstructed dendrite segment provides realistic spine geometries. a) Dendritic segment reconstructed from EM data stacks in [69]. We conduct simulations for the bistable model in all the numbered spines, shown below the segment. PSDs are denoted by black lines on the dendritic segment. b) Temporal dynamics of bound AMPAR at the top of a PSD for the 7 different realistic spines.

**Table S1:** Species in the cytoplasm

| Species | Initial condition [$\mu M$] | Diffusion rate [$\frac{\mu m^2}{s}$] | Ref |
| --- | --- | --- | --- |
| $Ca_{cyto}$ | 0.1 | 220 | [43] |
| CaM | 10 | 10 | [58] |
| CaCaM | 0 | 10 | [58] |
| Ng | 20 | 10 | [58] |
| CaMNg | 0 | 10 | [58] |
| CaMKIIp | 0 | 0.1 | [58] |
| CaMKII | 20 | 0.1 | [58] |
| CaN | 0 | 10 | [58] |
| CaNp | 1 | 10 | [58] |
| I1 | 0 | 10 | [58] |
| I1p | 1.8 | 10 | [58] |
| PP1 | 0 | 10 | [58] |
| PP1p | 0.27 | 10 | [58] |
| Aint* | 0.2628 | 0.45 | [7] |

\* Simulations were started with these initial conditions and run to steady state. See Table S3 and Table S12 for individual simulation steady states.

**Table S2:** Species on the membrane

| Species | Initial condition [$\frac{molecules}{\mu m^2}$] | Diffusion rate [$\frac{\mu m^2}{s}$] | Ref |
| --- | --- | --- | --- |
| Amem* | 12 | 0.45 | [11, 50] |
| PSD95* | 4000 | 0.015 | [18, 85] |
| ABound* | 1000 | 0.0045 | [11] |

\* Simulations were started with these initial conditions and run to steady state. See Table S3 and Table S12 for individual simulation steady states.

**Table S3:** Idealized Spine initial conditions for different spine volumes

| Spine type | Aint [$\mu M$] | Amem [$\#/\mu m^2$] | PSD95 [$\#/\mu m^2$] | Bound AMPAR [$\#/\mu m^2$] |
| --- | --- | --- | --- | --- |
| Thin 0.5 | 0.1725 | 7.875 | 3922.55 | 1077.45 |
| Thin 0.75 | 0.1735 | 7.92 | 3918 | 1082.025 |
| Thin 1 | 0.174 | 7.95 | 3914.5 | 1085 |
| Thin 1.25 | 0.17485 | 7.98 | 3911.5 | 1088.5 |
| Thin 1.5 | 0.1755 | 8.01 | 3908.1 | 1091.9 |
| Mushroom 0.5 | 0.17045 | 7.78 | 3933 | 1067 |
| Mushroom 0.75 | 0.1715 | 7.83 | 3927.8 | 1072.2 |
| Mushroom 1 | 0.17233 | 7.865 | 3923.8 | 1076.2 |
| Mushroom 1.25 | 0.173 | 7.9 | 3920.4 | 1079.6 |
| Mushroom 1.5 | 0.1736 | 7.925 | 3917.5 | 1082.45 |

**Table S4:**
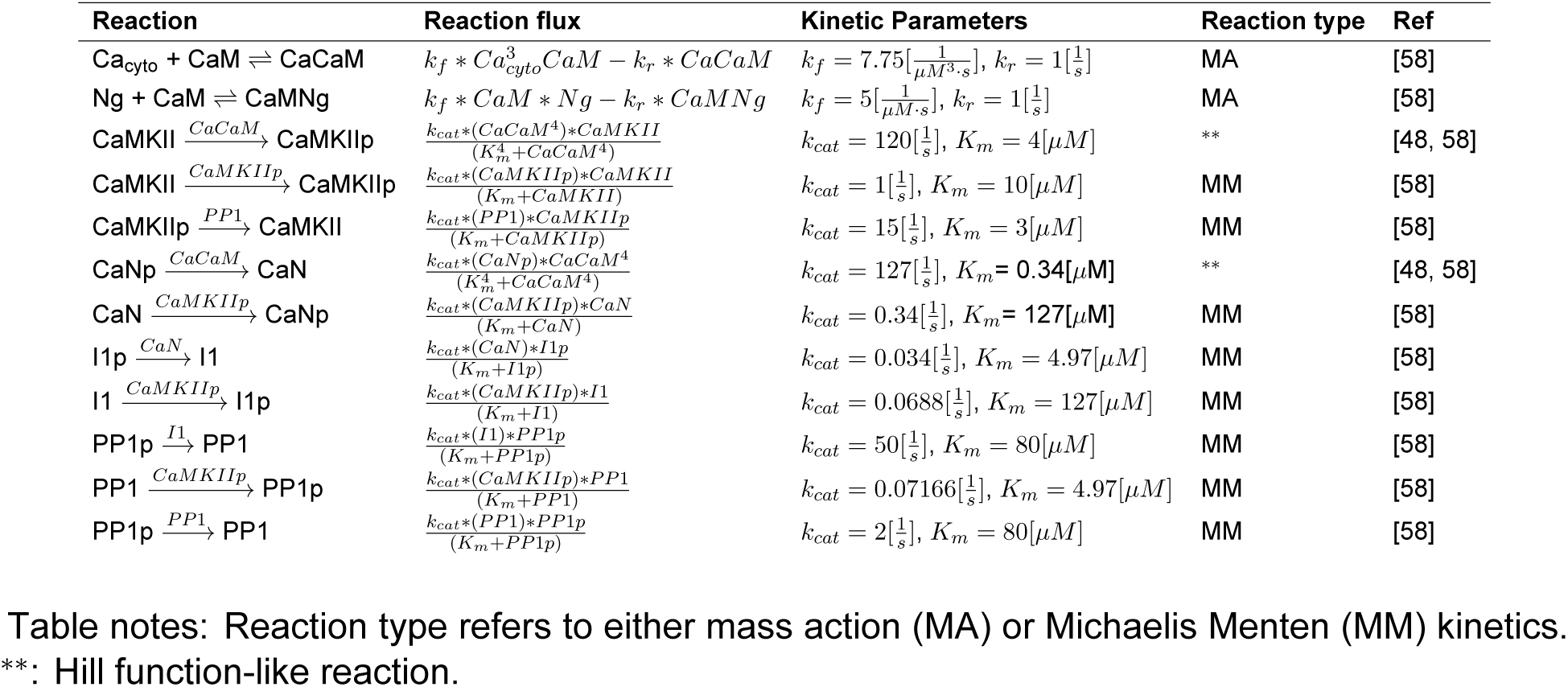
Cytosolic Reactions for bistable system

| Reaction | Reaction flux | Kinetic Parameters | Reaction type | Ref |
| --- | --- | --- | --- | --- |
| $\text{Ca}_{\text{cyto}} + \text{CaM} \rightleftharpoons \text{CaCaM}$ | $k_f * \text{Ca}_{\text{cyto}}^3 \text{CaM} - k_r * \text{CaCaM}$ | $k_f = 7.75[\frac{1}{\mu\text{M}^3 \cdot \text{s}}], k_r = 1[\frac{1}{\text{s}}]$ | MA | [58] |
| $\text{Ng} + \text{CaM} \rightleftharpoons \text{CaMNg}$ | $k_f * \text{CaM} * \text{Ng} - k_r * \text{CaMNg}$ | $k_f = 5[\frac{1}{\mu\text{M} \cdot \text{s}}], k_r = 1[\frac{1}{\text{s}}]$ | MA | [58] |
| $\text{CaMKII} \xrightarrow{\text{CaCaM}} \text{CaMKIIP}$ | $\frac{k_{\text{cat}} * (\text{CaCaM}^4) * \text{CaMKII}}{(K_m^4 + \text{CaCaM}^4)}$ | $k_{\text{cat}} = 120[\frac{1}{\text{s}}], K_m = 4[\mu\text{M}]$ | ** | [48, 58] |
| $\text{CaMKII} \xrightarrow{\text{CaMKIIP}} \text{CaMKIIP}$ | $\frac{k_{\text{cat}} * (\text{CaMKIIP}) * \text{CaMKII}}{(K_m + \text{CaMKIIP})}$ | $k_{\text{cat}} = 1[\frac{1}{\text{s}}], K_m = 10[\mu\text{M}]$ | MM | [58] |
| $\text{CaMKIIP} \xrightarrow{\text{PP1}} \text{CaMKII}$ | $\frac{k_{\text{cat}} * (\text{PP1}) * \text{CaMKIIP}}{(K_m + \text{CaMKIIP})}$ | $k_{\text{cat}} = 15[\frac{1}{\text{s}}], K_m = 3[\mu\text{M}]$ | MM | [58] |
| $\text{CaNp} \xrightarrow{\text{CaCaM}} \text{CaN}$ | $\frac{k_{\text{cat}} * (\text{CaNp}) * \text{CaCaM}^4}{(K_m^4 + \text{CaCaM}^4)}$ | $k_{\text{cat}} = 127[\frac{1}{\text{s}}], K_m = 0.34[\mu\text{M}]$ | ** | [48, 58] |
| $\text{CaN} \xrightarrow{\text{CaMKIIP}} \text{CaNp}$ | $\frac{k_{\text{cat}} * (\text{CaMKIIP}) * \text{CaN}}{(K_m + \text{CaN})}$ | $k_{\text{cat}} = 0.34[\frac{1}{\text{s}}], K_m = 127[\mu\text{M}]$ | MM | [58] |
| $\text{I1p} \xrightarrow{\text{CaN}} \text{I1}$ | $\frac{k_{\text{cat}} * (\text{CaN}) * \text{I1p}}{(K_m + \text{I1p})}$ | $k_{\text{cat}} = 0.034[\frac{1}{\text{s}}], K_m = 4.97[\mu\text{M}]$ | MM | [58] |
| $\text{I1} \xrightarrow{\text{CaMKIIP}} \text{I1p}$ | $\frac{k_{\text{cat}} * (\text{CaMKIIP}) * \text{I1}}{(K_m + \text{I1})}$ | $k_{\text{cat}} = 0.0688[\frac{1}{\text{s}}], K_m = 127[\mu\text{M}]$ | MM | [58] |
| $\text{PP1p} \xrightarrow{\text{I1}} \text{PP1}$ | $\frac{k_{\text{cat}} * (\text{I1}) * \text{PP1p}}{(K_m + \text{PP1p})}$ | $k_{\text{cat}} = 50[\frac{1}{\text{s}}], K_m = 80[\mu\text{M}]$ | MM | [58] |
| $\text{PP1} \xrightarrow{\text{CaMKIIP}} \text{PP1p}$ | $\frac{k_{\text{cat}} * (\text{CaMKIIP}) * \text{PP1}}{(K_m + \text{PP1})}$ | $k_{\text{cat}} = 0.07166[\frac{1}{\text{s}}], K_m = 4.97[\mu\text{M}]$ | MM | [58] |
| $\text{PP1p} \xrightarrow{\text{PP1}} \text{PP1}$ | $\frac{k_{\text{cat}} * (\text{PP1}) * \text{PP1p}}{(K_m + \text{PP1p})}$ | $k_{\text{cat}} = 2[\frac{1}{\text{s}}], K_m = 80[\mu\text{M}]$ | MM | [58] |

Table notes: Reaction type refers to either mass action (MA) or Michaelis Menten (MM) kinetics. \*\*: Hill function-like reaction.

**Table S5:**
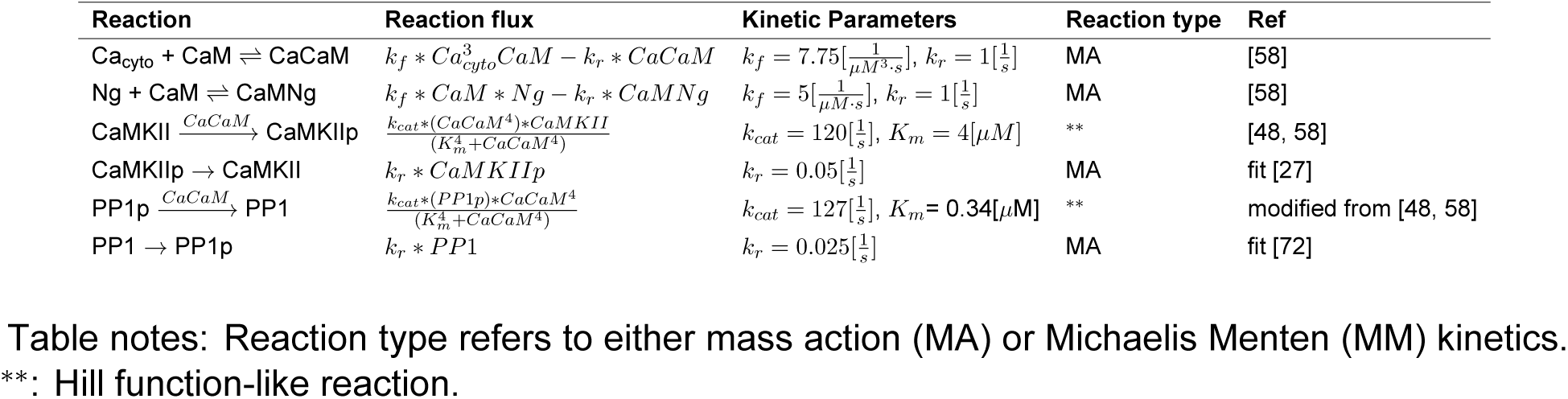
Cytosolic Reactions for monostable system

| Reaction | Reaction flux | Kinetic Parameters | Reaction type | Ref |
| --- | --- | --- | --- | --- |
| $\text{Ca}_{\text{cyto}} + \text{CaM} \rightleftharpoons \text{CaCaM}$ | $k_f * \text{Ca}_{\text{cyto}}^3 \text{CaM} - k_r * \text{CaCaM}$ | $k_f = 7.75[\frac{1}{\mu\text{M}^3 \cdot \text{s}}], k_r = 1[\frac{1}{\text{s}}]$ | MA | [58] |
| $\text{Ng} + \text{CaM} \rightleftharpoons \text{CaMNg}$ | $k_f * \text{CaM} * \text{Ng} - k_r * \text{CaMNg}$ | $k_f = 5[\frac{1}{\mu\text{M} \cdot \text{s}}], k_r = 1[\frac{1}{\text{s}}]$ | MA | [58] |
| $\text{CaMKII} \xrightarrow{\text{CaCaM}} \text{CaMKIIP}$ | $\frac{k_{\text{cat}} * (\text{CaCaM}^4) * \text{CaMKII}}{(K_m^4 + \text{CaCaM}^4)}$ | $k_{\text{cat}} = 120[\frac{1}{\text{s}}], K_m = 4[\mu\text{M}]$ | ** | [48, 58] |
| $\text{CaMKIIP} \rightarrow \text{CaMKII}$ | $k_r * \text{CaMKIIP}$ | $k_r = 0.05[\frac{1}{\text{s}}]$ | MA | fit [27] |
| $\text{PP1p} \xrightarrow{\text{CaCaM}} \text{PP1}$ | $\frac{k_{\text{cat}} * (\text{PP1p}) * \text{CaCaM}^4}{(K_m^4 + \text{CaCaM}^4)}$ | $k_{\text{cat}} = 127[\frac{1}{\text{s}}], K_m = 0.34[\mu\text{M}]$ | ** | modified from [48, 58] |
| $\text{PP1} \rightarrow \text{PP1p}$ | $k_r * \text{PP1}$ | $k_r = 0.025[\frac{1}{\text{s}}]$ | MA | fit [72] |

Table notes: Reaction type refers to either mass action (MA) or Michaelis Menten (MM) kinetics. \*\*: Hill function-like reaction.

**Table S6:**
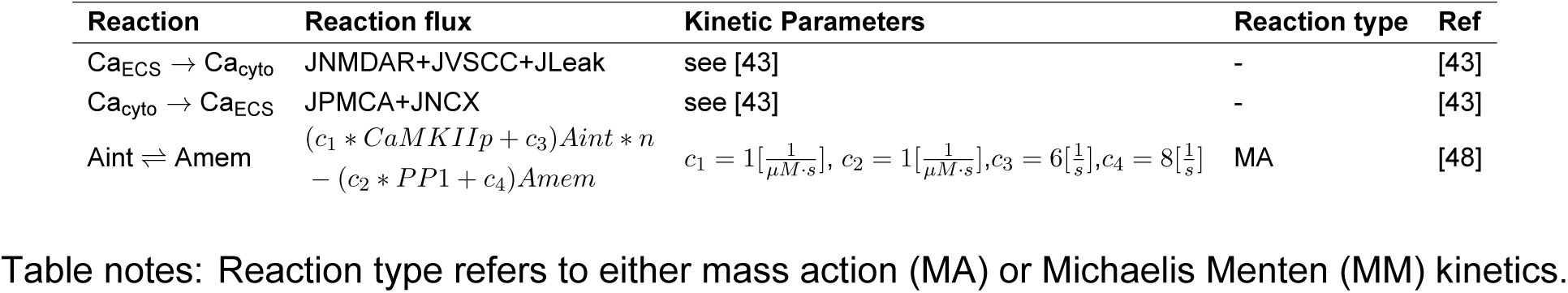
Membrane Flux Reactions

| Reaction | Reaction flux | Kinetic Parameters | Reaction type | Ref |
| --- | --- | --- | --- | --- |
| $\text{Ca}_{\text{ECS}} \rightarrow \text{Ca}_{\text{cyto}}$ | JNMDAR+JVSCC+JLeak | see [43] | - | [43] |
| $\text{Ca}_{\text{cyto}} \rightarrow \text{Ca}_{\text{ECS}}$ | JPMCA+JNCX | see [43] | - | [43] |
| $\text{Aint} \rightleftharpoons \text{Amem}$ | $(c_1 * \text{CaMKIIP} + c_3) \text{Aint} * n$<br>$- (c_2 * \text{PP1} + c_4) \text{Amem}$ | $c_1 = 1[\frac{1}{\mu\text{M} \cdot \text{s}}], c_2 = 1[\frac{1}{\mu\text{M} \cdot \text{s}}], c_3 = 6[\frac{1}{\text{s}}], c_4 = 8[\frac{1}{\text{s}}]$ | MA | [48] |

Table notes: Reaction type refers to either mass action (MA) or Michaelis Menten (MM) kinetics.

**Table S7:** Membrane surface reactions

| Reaction | Reaction flux | Kinetic Parameters | Reaction type | Ref |
| --- | --- | --- | --- | --- |
| $Aint \rightleftharpoons Amem$ | $(c_1 * CaMKIIp + c_3)Aint * n - (c_2 * PP1 + c_4)Amem$ | $c_1 = 1[\frac{1}{\mu M \cdot s}], c_2 = 1[\frac{1}{\mu M \cdot s}], c_3 = 6[\frac{1}{s}], c_4 = 8[\frac{1}{s}]$ | MA | [58] |
| $PSD95 + Amem \rightleftharpoons ABound$ | $k_f * PSD95 * Amem - k_r * ABound$ | $k_f = 0.0349[\frac{\mu m^2}{molecules \cdot s}], k_r = 1[\frac{1}{s}]$ | MA | this work |

Table notes: Reaction type refers to either mass action (MA) or Michaelis Menten (MM) kinetics.

**Table S8:** Membrane Surface Boundary Flux

| Reaction | Reaction flux | Kinetic Parameters | Ref |
| --- | --- | --- | --- |
| $A_{pool} \rightarrow Amem$ | $A_{base}/\tau_{base} \exp(-t/\tau_{decay}) * [CaMKIIp]$ | $A_{base}^* = 0.06546 [\mu m^2], \tau_{base} = 800 [s], \tau_{decay} = 60 [s]$ | this work, inspired by [22] |

\* Area of the spine base was changed based on spine geometry and therefore varied for each spine volume.

**Table S9:** Trafficking variations in terms of mathematical terms

| Case | Description |
| --- | --- |
| No endo/exo | $r_{mem} = 0$ |
| No pool | $A_{pool} = 0$ |
| No membrane diffusion | $D_{Am} = 0$ |

**Table S10:** Idealized Spine geometries

| Spine type | Volume [$\mu m^3$] | PM Area [$\mu m^2$] | PSD Area [$\mu m^2$] | Base Area [$\mu m^2$] |
| --- | --- | --- | --- | --- |
| Thin 0.5 | 0.020575 | 0.43873 | 0.025332 | 0.041 |
| Thin 0.75 | 0.030863 | 0.57489 | 0.033195 | 0.054 |
| Thin 1 | 0.040655 | 0.69644 | 0.040212 | 0.065 |
| Thin 1.25 | 0.051438 | 0.80814 | 0.046662 | 0.076 |
| Thin 1.5 | 0.061726 | 0.91259 | 0.051647 | 0.086 |
| Mushroom 0.5 | 0.13869 | 1.6364 | 0.19236 | 0.073 |
| Mushroom 0.75 | 0.20803 | 2.1443 | 0.25206 | 0.096 |
| Mushroom 1 | 0.27738 | 2.5976 | 0.30535 | 0.116 |
| Mushroom 1.25 | 0.34672 | 3.0142 | 0.35432 | 0.135 |
| Mushroom 1.5 | 0.41607 | 3.4038 | 0.40012 | 0.153 |

**Table S11:** Real Spine Geometries

| Spine type | Volume [$\mu m^3$] | PM Area [$\mu m^2$] | PSD Area [$\mu m^2$] | Base Area [$\mu m^2$] |
| --- | --- | --- | --- | --- |
| 1 | 0.091617 | 1.6743 | 0.071662 | 0.0386 |
| 2 | 0.11693 | 2.0386 | 0.033992 | 0.0295 |
| 3 | 0.041183 | 0.94775 | 0.041607 | 0.0365 |
| 4 | 0.075234 | 1.7054 | 0.02253 | 0.0354 |
| 5 | 0.025315 | 0.57447 | 0.040337 | 0.0142 |
| 6 | 0.46798 | 5.4388 | 0.41509 | 0.0451 |
| 7 | 0.23961 | 3.2347 | 0.13887 | 0.015 |

**Table S12:**
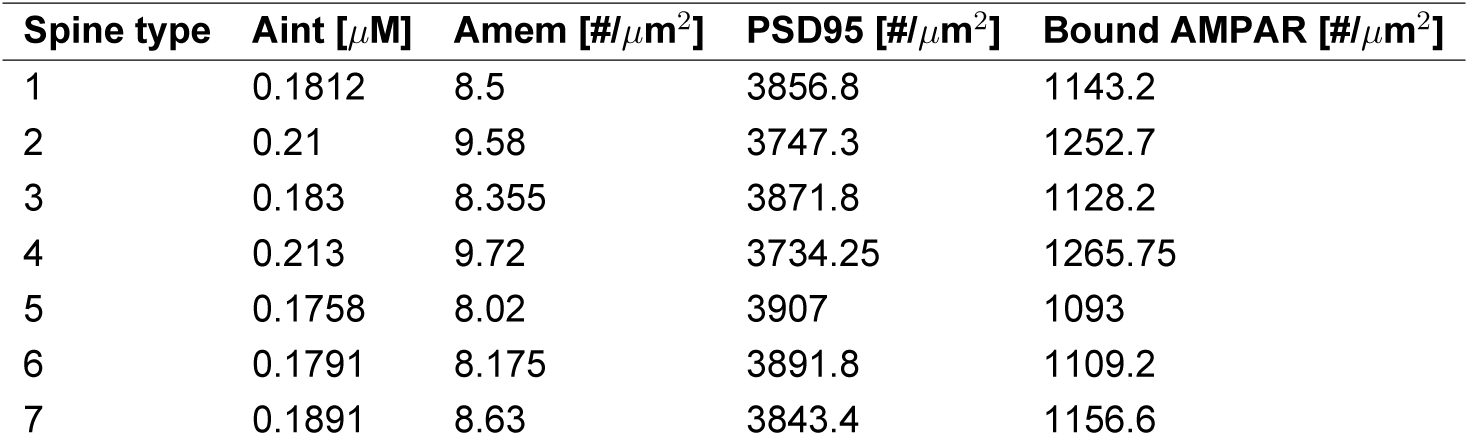
Realistic Spine initial conditions

| Spine type | Aint [$\mu M$] | Amem [$\#/\mu m^2$] | PSD95 [$\#/\mu m^2$] | Bound AMPAR [$\#/\mu m^2$] |
| --- | --- | --- | --- | --- |
| 1 | 0.1812 | 8.5 | 3856.8 | 1143.2 |
| 2 | 0.21 | 9.58 | 3747.3 | 1252.7 |
| 3 | 0.183 | 8.355 | 3871.8 | 1128.2 |
| 4 | 0.213 | 9.72 | 3734.25 | 1265.75 |
| 5 | 0.1758 | 8.02 | 3907 | 1093 |
| 6 | 0.1791 | 8.175 | 3891.8 | 1109.2 |
| 7 | 0.1891 | 8.63 | 3843.4 | 1156.6 |

